# Geography and forest naturalness as drivers of genetic diversity in saproxylic beetles

**DOI:** 10.1101/2025.11.29.691121

**Authors:** Nermeen R. Amer, Rama Sarvani Krovi, Alicja Wierzbicka, Radosław Plewa, Marcin Kadej, Tomasz Jaworski, Adrian Smolis, Andrzej Oleksa, Tomasz Szmatoła, Maria Oczkowicz, Łukasz Kajtoch

## Abstract

Understanding how geography and forest management shape genetic diversity is pivotal for the conservation of saproxylic beetles, the key indicators of forest ecosystem integrity. Using a multi-species Double Digest Restriction-site Associated DNA sequencing approach, we analyzed genome-scale SNP variation in 12 saproxylic beetle species (rare and common, incl. pests) differing in abundance, ecology, and mobility across central European forests, with respect to both geographic location and forest status. Analyses of distance-based redundancy test, molecular variance, and population structure consistently revealed that geography is the main determinant of genetic differentiation, with clear east-west clustering in several species. Forest status represented in four levels (commercial not protected, nature reserve, primeval not protected, and primeval protected) exerted a secondary but detectable effect in a subset of taxa, especially among rare and habitat-specialized species. For these species, populations from primeval protected sites showed distinct genetic structure compared to those from commercial stands. Our results highlight that regional connectivity, maintaining gene flow across a large spatial scale, should be prioritized in conservation planning, complemented by the protection of primeval forest refugia and the maintenance of structural complexity in commercial forests. By integrating high-resolution genomic data with geographical and management context, this study provides actionable insights into the processes shaping genetic structure in forest-dependent insects and underscores the importance of incorporating genetic monitoring into sustainable forest biodiversity management.

## Introduction

Forests are among the most extensive terrestrial ecosystems on the planet (Feng et al. 2016; Mori et al. 2017; Watson et al. 2018). Historical processes (such as ancient forest origins and post-glacial Holocene dispersal) have played a significant role in shaping the diversity, structure, and ecological heterogeneity of forested habitats (Ulrich 1995; Giesecke and Brewer 2018). Forests are widely recognized as biodiversity-rich systems, and a variety of global hotspots of species richness are located within forested regions (Gillespie et al. 2012; Bohn and Huth 2017; Mori et al. 2017), with profound implications, especially in areas under strong anthropogenic pressure. Trees, as primary biomass producers in forests, store energy in woody tissues such as xylem and phloem, making wood a vital resource for both natural communities and human use. In primeval forests, the diversity of organisms associated with wood is typically high; however, this diversity is significantly reduced in intensively managed forests, where only a limited number of species thrive (Christensen and Emborg 1996; Kuuluvainen 2009; Paillet et al. 2010a). While some organisms inhabit living trees, many are specialized to colonize dying and dead trees, often referred to as deadwood. For years, this ecological niche and its inhabitants were neglected in forest studies (Siitonen 2001), but growing awareness has led to increased research on wood-dependent taxa (Chave et al. 2006; Ogle et al. 2014; Kozák et al. 2021). This research is crucial as the functioning and dynamics of forest ecosystems are influenced by a variety of abiotic and biotic factors, including human activity (Guo et al. 2013; Rasche et al. 2013; Bayat et al. 2021).

The current diversity of wood-dwelling organisms is the result of past environmental and climatic conditions and recent changes caused by human activity. Forest development changed along Pleistocene glacial cycles. In Europe that forced forest-dwelling organisms to contract their ranges to the southernmost refugia or to moved eastward into Asia, while a minority survived in local cryptic refugia in Central or Western Europe (Taberlet et al. 1998; Hewitt 2000; Schmitt 2007). These shifts in species ranges had consequences for their biogeography (the distribution of distinct lineages) with important transient zones identified among many species or subspecies in Central Europe.

Later, in late Holocene, human activity, such as forest fragmentation, deforestation, and intensive forest management, has dramatically altered woodland ecosystems (Abdullah and Nakagoshi 2007; Taylor et al. 2009; Freer-Smith et al. 2019). This has especially impacted species dependent on old-growth microhabitats like deadwood (usually called saproxylic, although in this group should be listed not only xylophages or saprophages, but also mycetophages, and predators living in wood of dead or dying trees). Among saproxylic organisms, the most diverse (likely, except for microorganisms, which are poorly known) are arboreal fungi and insects, which are threatened by the depletion of such habitats in commercial woodlands (De Vivo et al. 2023). While some generalist species have acclimated to managed environments and even become pests, specialized species often experience reduced populations or local extinctions. These changes are now increasingly intensified by climate change, which continues to affect biodiversity and forest composition (Pawson et al. 2013; Thompson et al. 2014; Murray et al. 2017).

The most diverse and best known among saproxylic organisms are beetles (Coleoptera) (Hjältén et al. 2012; Thorn et al. 2018). Saproxylic beetle species that depend on dead trees, decaying wood, or associated fungi for at least part of their life cycle are crucial for forest ecosystems. They contribute to the decomposition of woody debris, nutrient cycling, serve as prey for a wide range of animals, or are predators on other saproxylic organisms (Grove 2002; Gimmel and Ferro 2018). Due to their robust association with deadwood habitats, many saproxylic beetles are highly sensitive indicators of forest biodiversity and ecological continuity (Hjältén et al. 2012). However, forest fragmentation and intensive logging, including sanitary cuttings, have led to the loss of deadwood microhabitats and extensive declines in saproxylic beetle populations (de Groot et al. 2019; Mazur et al. 2021). On the other hand, a large number of cambio- and xylophagous beetles (e.g., bark beetles, jevel beetles, longhorn beetles, etc.) are prone to outbreaks in commercial (planted, monospecific and even-aged) forests and are treated as pests in timber production, although they are still natural elements in old-growth forests (Singh et al. 2024).

Conservation or management efforts for these beetles require extensive knowledge of their population structure, genetic diversity, and gene flow. Traditional ecological and morphological assessments are often insufficient in resolving population-level patterns due to the frequent presence of cryptic taxa and the limited resolution power of these methods. In contrast, molecular methods provide powerful tools for determining genetic variability and connectivity among populations, which are vital for understanding their resilience, enhancing conservation strategies, and effective management (Audisio et al. 2009; Eberle et al. 2021; Krovi et al. 2024). Recently, knowledge on population genetics of saproxylic beetles had been summarized (Kajtoch et al. 2019; Krovi et al. 2024) supporting the their complex genetic variability of across Europe, with Central European forests as a transient zone between various evolutionary units and a crucial region for understanding the evolution and ecology of deadwood-dwelling species. Although all previous studies were done on traditional molecular markers (e.g. mitochondrial DNA or microsatellites), whereas genomic data started to be implemented for saproxylic beetles in Europe very recently.

High-throughput sequencing (HTS) techniques, like genotyping-by-sequencing (GBS), double-digest restriction-site associated DNA sequencing (ddRADseq), or whole genome sequencing (WGS), and their variants, have revolutionized population genetics by providing cost-effective, genome-wide discovery of single-nucleotide polymorphisms (SNPs) in non-model organisms (Peterson et al. 2012). Among these, ddRADseq has become a commonly used method due to its high reproducibility, customizable coverage, and low error rates (Burns et al. 2017; Aguirre et al. 2019). ddRADseq is particularly suitable for exploring population genetics in diverse taxa with little or no prior genetic resources (Peterson et al. 2012; Tóth et al. 2021; Magbanua et al. 2023).

SNP markers have emerged as powerful molecular markers for conservation genetics, assessing genetic diversity and delineating conservation units in animal populations (Vignal et al. 2002; Coates et al. 2011; Wenne 2023). Their abundance and genome-wide distribution make them valuable for non-model organisms, including many beetle species. For example, thousands of high-quality SNPs supported three beetle taxa (*Lucanus* spp.), SNPs developed through the ddRAD technique enhanced the analysis of genetic diversity, yielding clear management implications of each lineage (De Vivo et al. 2023). SNPs generated from WGSs proved powerful for detecting the demographic history of bark beetles (*Ips typographus*) in Europe (Ellerstrand et al. 2022). Hence, SNPs play a vital role in the conservation genetics of rare taxa or in the population genetics of pests, providing high-resolution insights into genetic diversity and population structure. Such detailed information facilitates the identification of the evolutionary significant units (ESUs) (Casacci et al. 2014) and management units (MUs) (Palsbøll et al. 2007), which supports the development of conservation strategies to maintain genetic variation and adaptive capacity within targeted populations (Fraser and Bernatchez 2001; Barbosa et al. 2018).

In this study, we used ddRADseq to assess the genetic diversity and population structure of several saproxylic beetle species in Central European forests. These species were sampled across a wide range of forest types in Poland, including primeval and commercial stands, both protected and non-protected. Therefore, our objectives are: (1) to quantify genetic diversity between beetle populations collected from forests localized in a presumed a transient zone in Central Europe to verify the hypothesis that westernmost and easternmost forests are inhabited by significantly differentiated populations; (2) to see if forest status (management or conservation) affect saproxylic beetles’ genetic diversity, that is to test the hypothesis that populations from primeval forests are more diverse than these in commercial ones); (3) to assess if observed patterns are consistent among various species (e.g. rare vs common); and (4) to provide data that can support conservation efforts by identifying populations with low genetic variability and/or high isolation. This research contributes to the growing fields of population and conservation genetics, highlighting the importance of integrating molecular tools into biodiversity monitoring programs, especially for ecologically significant but often overlooked taxa, such as saproxylic beetles.

## Material and methods

### Study Sites

Beetle samples were collected between 2022 and 2023 from eight forest complexes across Poland, spanning lowland, upland, and mountainous regions, and including forests representing the northernmost, southernmost, westernmost, and easternmost populations of selected species (Fig. S1, Table S1). Sites were chosen to demonstrate a gradient of forest types, from protected primeval forests (Białowieża F., Holy Cross Mountains, and the Carpathians), to commercial forests (Augustów F., Knyszyn F., Silesian F., Oder F., Barycz F.) (Fig. S1, Table S1). In the latter forests, beetles were collected from typical commercial stands (if present due to a shortage of deadwood resources), referred to as “commercial” throughout the manuscript, and from nature reserves (with old-growth forests resembling natural ones, although of restricted area and isolated in a commercial woodland matrix), referred to as “nature reserve” throughout the manuscript. Also in primeval forests, beetles were collected from both protected stands (national parks and nature reserves) referred to as “primeval protected”, and managed stands (although of much lower logging intensity than in remaining management forests) referred to as “primeval not protected” throughout the manuscript. To simplify some analyses, samples from following forests were grouped, accordingly to geographic location of a given forests: NE (Białowieża F., Knyszyn F., Augustów F. – all from Podlasie lowlands); SW (Silesian F., Oder F., Barycz F. – all from Silesia lowlands); and SE (Holy Cross Mountains, and the Carpathians – both from mountains). Finally, thousands of sites were visited, but only 421 sites hosted selected beetles, with some sites hosting more than one taxon (Table S1).

### Sample Collection

In each forest complex, beetles were searched for using various techniques: i) inspection of microhabitats under the bark of dying or dead trees, ii) cutting of decayed deadwood, iii) inspection of fungi (also using a sieve), iv) funnel and barrier traps with pheromone lures for selected taxa.

Sampling focused on ecologically important saproxylic beetle species representing various trophic guilds (saprophages, xylophages, mycetophages, predators) and having different abundance or status (rare/threatened and protected, common, or pest) (Table 1). In this study, species that are frequently present across all sites are referred to as “common” throughout the manuscript. A subset of these species that are known to cause damage to deadwood during population outbreaks are referred to as “pests”, while species occurring infrequently or in low numbers are referred to as “rare” throughout the manuscript. Finally, among selected beetles, 12 species were selected based on the above-listed criteria and sampling availability (for which at least several sites were available across all examined forests). Among selected rare taxa were relics of primeval forests (Eckelt et al. 2018) (*Boros schneideri, Cucujus cinnaberinus, Neomida haemorrhoidalis,* and *Elater ferrugineu*s), common taxa (e.g. *Thanasimus formicarius*, *Bolitophagus reticulatus, Schizotus pectinicornis, Stenurella melanura*), and pests (e.g. *Ips acuminatus, Ips typographus, Rhagium inquisitor,* and *Phaenops cyanea*). *Rhagium inquisitor* is not causing death of trees (like other listed species in this group), but its larvae damage wood, therefore it is considered as pest, at least in some regions. The list of these species, along with their characteristics, is presented in Table 1. Some of the selected species were collected as adults, whereas others were collected as larvae, as only this life stage could be found relatively easily in the field (Table 1). Detailed information on the sample sizes is provided in Table S2.

**Table 1.**
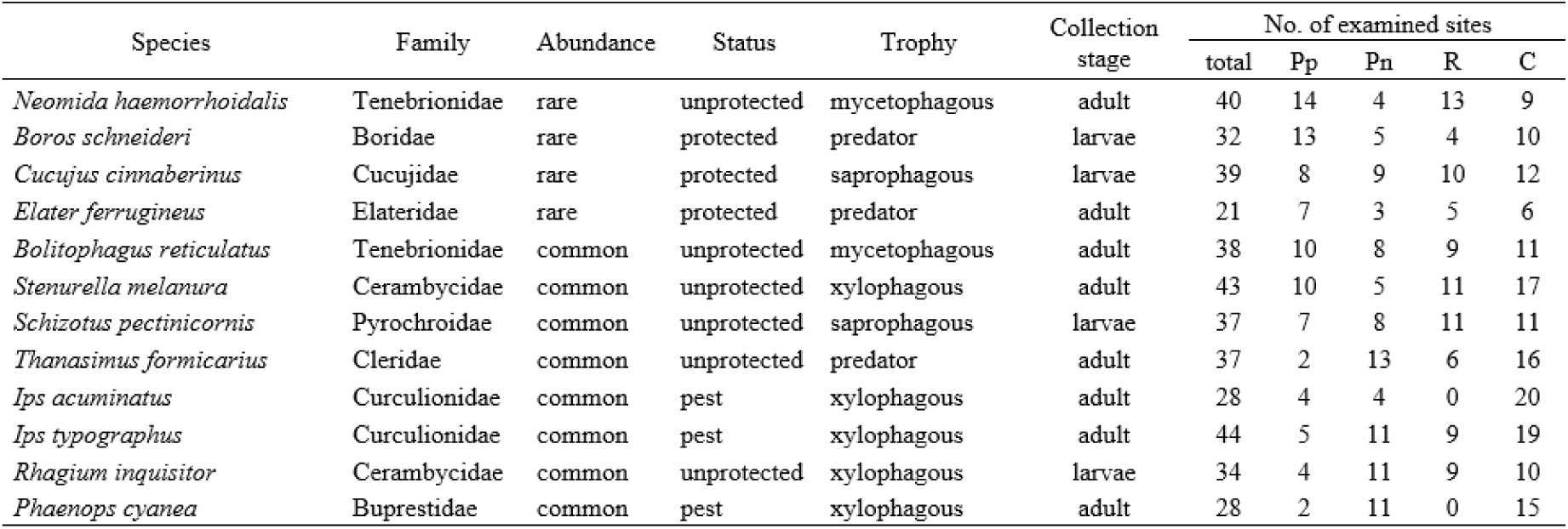
Summary of species taxonomic classification, ecological characteristics, sampling details, and number of examined sites (Pp: Primeval protected sites, Pn: Primeval not protected sites, R: Natural reserve sites, C: Commercial stand sites) used in this study.

Each specimen was stored in 95-99% ethanol and preserved at –20°C until DNA extraction. Each individual was determined to the species by specialists (coleopterologists) on the research team. No selected beetle taxa required the use of special techniques for identification (e.g., preparation of reproductive organs, etc.).

### Laboratory procedures

#### DNA Extraction

Prior to DNA isolation, each individual was cleaned with distilled water to remove external contaminants. Each individual was processed separately in sterile conditions.

For small beetles (<1 cm in length), the entire body was used for DNA extraction. In larger beetles, only the head with thorax, and legs were used, whereas the abdomen was left (stored). For large larvae, only the head and the first three segments were used for DNA isolation. For the largest species (*E. ferrugineus*), only legs were collected due to restrictions imposed by nature protection authorities.

DNA was extracted using Sherlock AX (A&A Biotechnology) according to the manufacturer’s protocols.

#### Library preparation and sequencing

ddRad-seq libraries were prepared according to the method described by Bayona-Vásquez et al. (2019). Briefly, DNA concentration was normalized to 20ng/μL and 5 μL of each sample was transferred to 96-well plate. Next, iTru Fusion Primers with Internal Tags (iTru_NheI_R1_stub_A – H and iTru_EcoRI_R2_RC_stub_1-12), and appropriate restriction enzymes were added to identify each sample within the plate. After 1 hour incubation at 37 °C, ligation was performed with T4 ligase. Ligation products were pooled (96 samples into two tubes of 240 μL) and, after purification with AMPure XP magnetic beads (Beckman Coulter), PCR was performed with i5 and i7 primers to identify each pool during sequencing. PCR products were again purified with Agencourt AMPure XP beads and quantified using a Qubit. Finally, the library range was analyzed on the Tape station, and fragments between 400 and 600 bp were cut using the Pippin Prep instrument (Sage Science). Libraries from 4 plates (384 samples) were pooled, normalized to 2nM, and sent to the sequencing utility. Pooled libraries were sequenced at Admera Health Company (New Jersey, USA) on an Illumina NovaSeq X Plus - 10B - PE 150, yielding approximately 4 million reads per sample.

### Bioinformatic analyses

#### Quality Control of Raw Sequencing Data

Quality assessment of the raw FASTQ files was performed using FastQC v0.11.9 (Andrews 2010), a standard tool for checking the quality of high-throughput sequencing data. Since this is a ddRADseq dataset generated with 141bp paired-end reads, all sequences have a uniform length determined by the sequencing protocol. Therefore, no explicit minimum read length filter was applied at this stage. Each sequencing file was assessed individually for key metrics, including per-base sequence quality, GC content, sequence length distribution, adapter contamination, and duplication levels. Summarized quality reports were generated and reviewed using MultiQC v1.32 (Ewels et al. 2016) to identify samples with abnormal patterns or quality concerns. Reads with low-quality scores (Phred < 30) or uncalled bases were removed. Only reads that passed quality filtering and retained their full restriction-site overhangs were retained for further processing using the Stacks pipeline. This rigorous quality control ensured high-confidence SNP calling and minimized downstream biases in population genetic analyses.

#### Alignment and variant calling (Stacks Pipeline)

Raw reads were demultiplexed, quality-filtered, and trimmed using the process_radtags module in Stacks v2.68 (Rochette et al. 2019b). De novo locus assembly was performed using the ustacks, cstacks, and sstacks modules. Key parameters included a minimum stack depth (-m = 3) and a maximum number of mismatches (-M = 2) per individual. The catalog was built using cstacks across all individuals with a mismatch allowance (-n = 1) between loci.

Detection and genotyping of all the loci from the aligned reads were done by using the gstacks program included in Stacks v2.68. The detected loci were filtered using the software ‘populations’ to call SNPs and compute population-level statistics, which is included in Stacks v2.68. It was run with (–r = 0.80) to retain only the loci that are present in at least 80% of the population. The following filters were also used to ensure high-genotyping quality:

- Minimum minor allele frequency (MAF of 0.03).
- Exclusion of samples with an average coverage lower than 5X from further analysis (--min-gt-depth 5).
- Only the positions that were informative in at least 70% of the individuals were retained for further analysis (Geno of 0.3).

Output files were exported in VCF and STRUCTURE formats for downstream analyses.

### Population genetic and statistical analysis

#### Genetic data preparation for downstream analysis

All ddRADseq datasets were processed using the same SNP filtering pipeline applied to each species separately (MAF=0.03, GENO=0.3). The identical filtering criteria ensured methodological consistency across species. Differences in the final number of retained SNPs and loci among species (Table S1), reflect neutral variation in genome characteristics (e.g., heterozygosity, restriction site distribution, and sequence depth) rather than differences in data processing. To account for linkage disequilibrium, SNP datasets for each species separately were LD-pruned prior to ADMIXTURE analysis using a threshold of (r2 < 0.1). For other analyses (distance-based redundancy analysis-dbRDA, Analysis of molecular variance-AMOVA, Discriminant analysis of principal component-DAPC, and heatmaps), the full filtered SNP sets were used without LD pruning to retain the maximum genetic information, as these multivariate and distance-based approaches are less affected by linkage (Excoffier et al. 1992; Jombart et al. 2010). The number of SNPs retained after LD pruning for each species is provided in Table S2.

#### Spatial genetic patterns

We partitioned variance in pairwise genetic differentiation using distance-based redundancy analysis (dbRDA) with linearized F_st_ (F_st_/1-F_st_) as the response variable (Legendre and Andersson 1999). Management was encoded as a categorical predictor with four management levels (commercial, nature reserve, primeval-not-protected, and primeval-protected). Spatial structure was modeled using dbMEM eigenfunctions derived from site coordinates (Peres-Neto and Legendre 2010). For each species, we calculated pairwise Weir-Cockerham F_st_ among populations/sites and used the linearized F_st_ (F_st_ (F_st_/1-F_st_)) as the response variable to meet the distance-based model assumptions (Weir and Clark 1984; Rousset 1997). The geography predictor was calculated as distance (Km; log transformed), and the management predictor was estimated as categorical dummies for four levels. For each species, we quantified the unique variance explained by management (controlling for space) and by space (controlling for management) using partial R^2^. Significance p-value and partial R2 were obtained using a permutation-based model (permutations=9.999) for each predictor.

The AMOVA test (Excoffier et al. 1992) was quantified separately for populations grouped by management type and by geographic region. Variance was partitioned among and within populations separately for four management groups (commercial, nature reserve, primeval-not-protected, and primeval-protected), and three geography groups (North-East (NE), South-East (SE), and South-West (SW)). Significance was assessed by permutation tests (9,999 permutations) for each species separately.

For each species, to evaluate the relationship between genetic diversity and geographic distance under different management levels, we performed a distance decay analysis using pairwise linearized F_st_, which is merged with geographic coordinates and management categories (metadata file). Geographic distances (km) were calculated using the geosphere package (Hijmans et al. 2025). Isolation by distance and management effect for each species was quantified using multiple matrix regression with randomization (MMRR) implemented in the ecodist package (Goslee and Urban 2007), with 9.999 permutations. The response variable was linearized F_st_, and the predictors were the log-transformed geographic distance and management (four levels).

#### Population genetic structure analyses

Population structure was assessed using ADMIXTURE v.1.3 (Alexander and Lange 2011). The most probable number of ancestral clusters, K, is determined by cross-validation and low linkage disequilibrium (LD) values (< 0.1). For each individual, the admixture algorithm returns the probability Q (Q_1_, Q_2_, …, Q_10_) that it belongs to each of the K populations. The resulting individual ancestry coefficients (Q-matrix) were imported into R and visualized as stacked barplots with individuals ordered by populations based on four management levels (commercial, nature reserve, primeval-not-protected, and primeval-protected), and three geography groups (northeast (NE), southeast (SE), and southwest (SW)) using a metadata file.

Discriminant analysis of Principal components (DAPC) (Jombart et al. 2010) was implemented using the package adegenet (Jombart 2008). Genotypes were imported from a structure format file into a genind object, with individuals assigned to four management groups (commercial, nature reserve, primeval-not-protected, and primeval-protected) and three geography groups (northeast (NE), southeast (SE), and southwest (SW)). Individual scores in DA1 and DA2 were visualized with 95% normal data confidence. All plots were visualized using the ggplot2 package.

Pairwise F_st_ was quantified separately for populations grouped by management type and by geographic region. It was estimated for all population pairs with permutation-based p-value and BH-adjusted false discovery control; the resulting F_st_ matrix was visualized as a heatmap (rows/columns ordered by hierarchical clustering) to highlight clusters and gradients of differentiation.

For statistical analyses, we used R version 4.0.3 (R Development Core Team, 2013). All visualizations have been done using the ggplot2 package (Wickham 2016).

## Results

In general, our analyses indicated that spatial separation among populations is the predominant factor affecting genetic differentiation.

### Distance-based redundancy analysis (dbRDA)

Distance-based redundancy analysis (dbRDA) showed variable contributions of geography and management factors to genetic diversity among 12 beetle species (Table 2). Overall, dbRDA results suggest that geographic distance exerts a stronger influence on genetic differentiation between populations than forest management (Table 2). Geographic distance explained a more significant proportion of genetic differentiation in several species, with the highest R^2^ values in the common species (*S. melanura*, *S. pectinicornis*, *T. formicarius*, and *B. reticulatus*) followed by pests (*I. acuminatus*, *I. typographus*, and *P. cyanea*), indicating a clear spatial genetic structure consistent with isolation by distance (Fig.1, Table 2). Rare species showed little to no genetic differentiation, except for *N. haemorrhoidalis,* which showed a relatively moderate R^2^ proportion (R^2^=0.228). In contrast, management effects contributed less to overall variation, with low and mostly non-significant unique R^2^ values across all species, except for a weakly significant trend (p management = 0.09) in *R. inquisitor* and *P. cyanea* (Table 2).

**Fig. 1.**
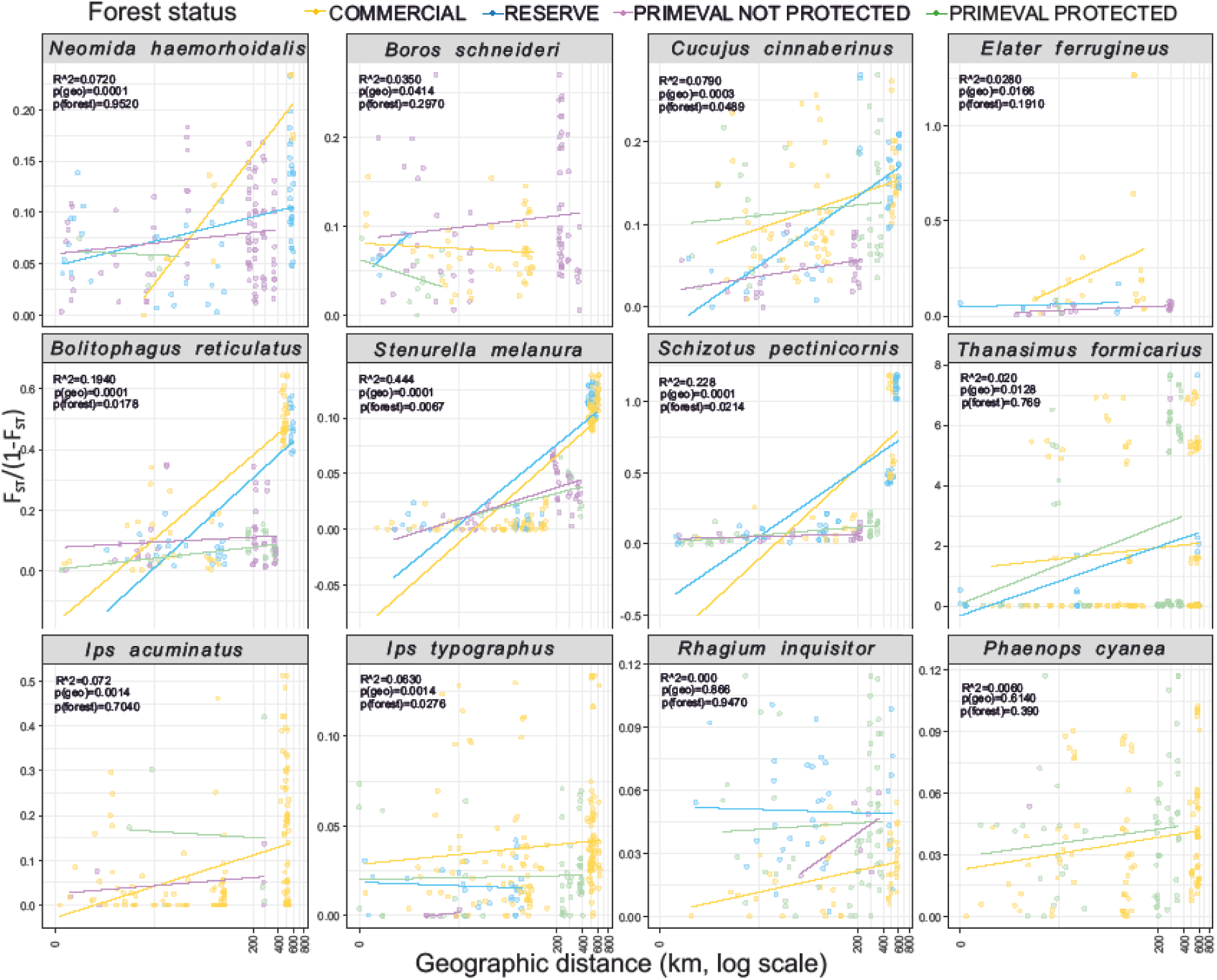
Isolation by distance (IBD) based on multiple matrix regression with randomization (MMRR) across the four forest management levels (commercial, nature reserve, primeval not protected, and primeval protected).

**Table 2.** Summary of distance-based redundancy analysis (dbRDA) results showing the unique and total variance in genetic diversity explained by geographic distance and forest management levels for 12 saproxylic beetle species. Reported are partial R2 values, permutation-based p-values (9.999 permutations) for each predictor (geography and management), and full R2 values for each species. The significance was determined using a threshold of P-value < 0.05.

| Species | No. sites | No. pairs | No. of SNPs | Geography levels | Managment levels | $R^2$ geography | P-value geography | $R^2$ management | P-value management | $R^2$ full |
| --- | --- | --- | --- | --- | --- | --- | --- | --- | --- | --- |
| *Neomida haemorrhoidalis* | 34 | 561 | 10501 | 3 | 4 | 0.228 | <0.001 | 0.007 | 0.518 | 0.264 |
| *Boros schneideri* | 31 | 465 | 6574 | 3 | 3 | 0.000 | 0.826 | 0.000 | 0.714 | 0.006 |
| *Cucujus cinnaberinus* | 39 | 741 | 2480 | 3 | 4 | 0.085 | <0.001 | 0.009 | 0.382 | 0.114 |
| *Elater ferrugineus* | 21 | 207 | 10210 | 3 | 4 | 0.000 | 0.555 | 0.000 | 0.872 | 0.000 |
| *Bolitophagus reticulatus* | 38 | 703 | 2841 | 3 | 4 | 0.398 | <0.001 | 0.234 | 0.346 | 0.486 |
| *Stenurella melanura* | 42 | 861 | 12063 | 3 | 4 | 0.707 | <0.001 | 0.011 | 0.956 | 0.786 |
| *Schizotus pectinicornis* | 35 | 595 | 2512 | 3 | 4 | 0.552 | <0.001 | 0.010 | 0.593 | 0.716 |
| *Thanasimus formicarius* | 37 | 666 | 2426 | 3 | 4 | 0.436 | <0.001 | 0.059 | 0.135 | 0.454 |
| *Ips acuminatus* | 26 | 325 | 3957 | 3 | 4 | 0.251 | 0.031 | 0.118 | 0.149 | 0.266 |
| *Ips typographus* | 41 | 820 | 1484 | 3 | 4 | 0.113 | 0.001 | 0.000 | 0.999 | 0.094 |
| *Rhagium inquisitor* | 34 | 561 | 4083 | 3 | 4 | 0.099 | <0.001 | 0.046 | 0.090 | 0.093 |
| *Phaenops cyanea* | 28 | 378 | 27583 | 3 | 3 | 0.079 | 0.097 | 0.088 | 0.09 | 0.125 |

### Genetic structure- AMOVA test-geography

AMOVA revealed greater genetic structure among populations categorized by geographic regions than among those categorized by forest management, with significant geographic structure in all species (Table 3). The percentages of variance between populations ranged from 1.5% to 50.7%, with the strongest differentiation shown in common taxa (e.g., *S. pectinicornis, T. formicarius, B. reticulatus, and S. melanura*) (Table 3). Intermediate differentiation was detected in rare taxa and was comparatively weak in a subset of pest species (except for *I. acuminatus*) (Table 3).

**Table 3.** Analysis of Molecular Variance (AMOVA) results for the studied 12 beetle species based on forests’ geography regions and management separately. The significance was determined using a threshold of P-value <0.05.

| Species | No. of ind. | No. of SNPs | df | Geography |  |  |  |  |  | Management |  |  |  |  |  |  |
| --- | --- | --- | --- | --- | --- | --- | --- | --- | --- | --- | --- | --- | --- | --- | --- | --- |
| | | | | $\sigma^2$ among | % among | $\sigma^2$ within | % within | P-value | $\Phi_{ST}$ | df | $\sigma^2$ among | % among | $\sigma^2$ within | % within | P-value | $\Phi_{ST}$ |
| *Neomida haemorrhoidalis* | 199 | 10501 | 2 | 14.371 | 5.814 | 233.150 | 94.191 | 0.0001 | 0.058 | 3 | 6.760 | 2.762 | 237.960 | 97.240 | 0.0001 | 0.027 |
| *Boros schneideri* | 158 | 6574 | 2 | 1.921 | 3.191 | 58.430 | 96.810 | 0.0001 | 0.032 | 3 | 1.563 | 2.621 | 58.231 | 97.381 | 0.0001 | 0.026 |
| *Cucujus cinnaberinus* | 190 | 2480 | 2 | 2.091 | 5.991 | 32.711 | 94.012 | 0.0001 | 0.059 | 3 | 1.241 | 3.631 | 33.011 | 96.372 | 0.0001 | 0.036 |
| *Elater ferrugineus* | 84 | 10210 | 2 | 45.342 | 8.173 | 509.570 | 91.821 | 0.0001 | 0.081 | 3 | 38.321 | 6.990 | 509.340 | 93.000 | 0.0001 | 0.069 |
| *Bolitophagus reticulatus* | 185 | 2841 | 2 | 35.571 | 23.421 | 116.280 | 76.572 | 0.0001 | 0.234 | 3 | 7.710 | 5.491 | 132.711 | 94.513 | 0.0001 | 0.054 |
| *Stenurella melanura* | 199 | 12063 | 2 | 43.592 | 12.311 | 310.543 | 87.681 | 0.0001 | 0.123 | 3 | 5.871 | 1.721 | 334.205 | 98.274 | 0.0001 | 0.017 |
| *Schizotus pectinicornis* | 167 | 2512 | 2 | 50.240 | 50.662 | 48.910 | 49.333 | 0.0001 | 0.506 | 3 | 10.941 | 13.073 | 72.770 | 86.921 | 0.0001 | 0.131 |
| *Thanasimus formicarius* | 178 | 2426 | 2 | 129.491 | 46.331 | 150.008 | 53.660 | 0.0001 | 0.046 | 3 | 23.351 | 9.940 | 211.532 | 90.053 | 0.0007 | 0.099 |
| *Ips acuminatus* | 125 | 3957 | 2 | 6.434 | 9.582 | 60.631 | 90.413 | 0.0001 | 0.095 | 2 | 1.221 | 1.891 | 63.232 | 98.120 | 0.051 | 0.018 |
| *Ips typographus* | 201 | 1484 | 2 | 0.411 | 4.180 | 9.442 | 95.823 | 0.0001 | 0.041 | 3 | 0.112 | 1.223 | 9.560 | 98.773 | 0.003 | 0.012 |
| *Rhagium inquisitor* | 164 | 4083 | 2 | 0.703 | 2.411 | 28.562 | 97.594 | 0.0001 | 0.02 | 3 | 0.311 | 1.070 | 28.731 | 98.931 | 0.0002 | 0.011 |
| *Phaenops cyanea* | 140 | 27583 | 2 | 15.660 | 1.463 | 1052.651 | 98.534 | 0.0002 | 0.014 | 2 | 11.272 | 1.060 | 1056.071 | 98.942 | 0.0006 | 0.011 |

### Genetic structure-AMOVA test-management

AMOVA partitioning by management showed that most genetic variation occurred within populations, while only a small proportion was explained by differences among management levels (Table 3). The percentages of variance between populations ranged from 1.2% to 13.1%, with the highest differentiation shown in the common species *S. pectinicornis* and *T. formicarius* (Table 3). Moderate differentiation was observed in the rare-threatened species *E. ferrugineus* and the common-habitat specific species *B. reticulatus* (Table 3). Only *I. acuminatus* showed a marginally non-significant result (p=0.05) (Table 3).

### Distance – decay MMRR

Across the 12 species, distance decay plots showed that genetic diversity generally increased with increasing geographic distance, with a significant geographic distance effect in all species except for *R. inquisitor* and *P. cyanea* (Fig. 1). Management effects were weaker and more idiosyncratic where unique management levels were significant in five species (three common: *S. melanura*, *S. pectinicornis*, and *B. reticulatus*, one pest: *I. typographus*, and one rare species: *C. cinnaberinus*).

### Genetic structure- DAPC- geography

Across the 12 species for which we ran DAPC by region (NE, SE, and SW), we observed heterogeneous but overall geography-driven structure (Fig. 2). DAPC showed strong genetic structure in common species (*S. pectinicornis*, *B. reticulatus*, and *S. melanura*), each forming well-separated (east-west) regional clusters. In contrast, the remaining common and pest taxa (*T. formicarius*, *I. typographus*, *I. acuminatus*, and *R. inquisitor*) exhibit broad overlap among regions, consistent with higher connectivity. The rare species: *C. cinnaberinus*, *N. haemorrhoidalis*, and *E. ferrugineus exhibit* intermediate genetic structure across populations. Some taxa (*R. inquisitor*, *E. ferrugineus*, and *I. typographus*) showed variable regional clustering separation (north-south) (Fig. 2). One species, *B. schneideri*, lacking a SW population, was replaced with another population to show the differences between NE and SE populations.

**Fig. 2.**
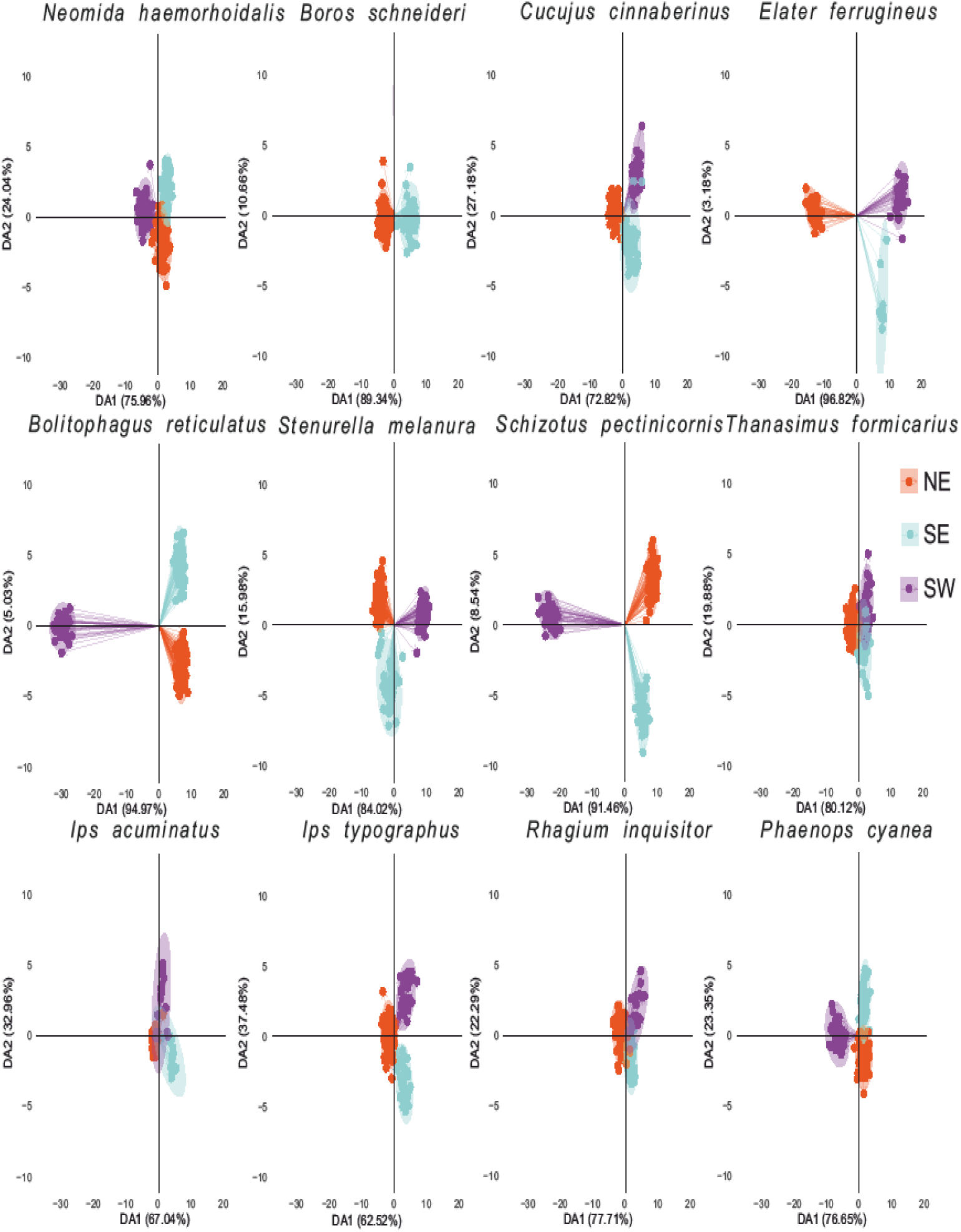
Genetic structure represented by discriminant analysis of principal components (DAPC) based on forests’ geographic regions (NE: north east, SE: south east, SW: south west).

### Genetic structure- DAPC- management

Across the 12 beetle species for which we ran DAPC by management level (commercial, nature reserve, primeval not protected, and primeval protected), we observed a moderate heterogeneous management-driven structure (Fig. 3). DAPC panels of species show the highest pronounced separation between the primeval protected and commercial forests in four species (*S. pectinicornis*, *B. reticulatus*, *E. ferrugineus*, and *C. cinnaberinus*), with partial weak overlapping in (rare species: *N. haemorrhoidalis*, *B. schneideri*, common: *S. melanura*, and pest: *P. cyanea*). In contrast, the remaining common/pest taxa (*T. formicarius*, *R. inquisitor*, *I. typographus*, and *I. acuminatus*) exhibit wide overlap among management levels (Fig. 3).

**Fig. 3.**
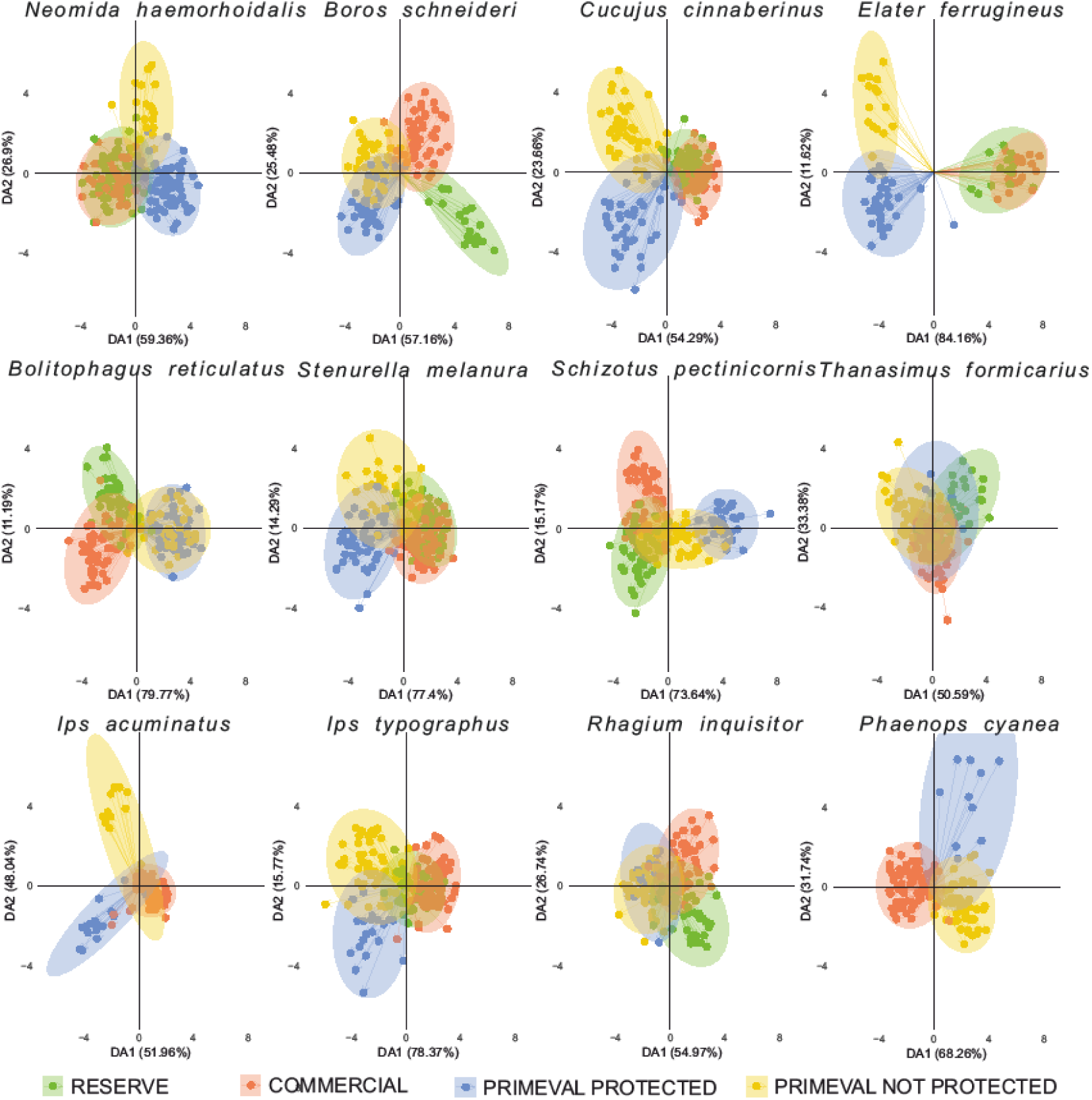
Genetic structure represented by discriminant analysis of principal components (DAPC) based on four forest management levels (commercial, nature reserve, primeval not protected, and primeval protected).

### Genetic structure- Admixture-geography

Ancestry assignments are consistent with DAPC ranking. The common species (*B. reticulatus and S. pectinicornis*), the pest (*I. acuminatus*), and the rare taxa (*N. haemorrhoidalis* and *E. ferrugineus*) display regional coherent ancestry blocks with limited cross-assignment among populations (NE, SE, SW) (Fig. 4). In contrast, the species (*S. melanura*, *T. formicarius*, *R. inquisitor*, *I. typographus*, *P. cyanea*, *C. cinnaberinus*, and *B. schneideri* show more mixed ancestry across regions, indicating weaker geographic partitioning of gene pools (Fig. 4).

**Fig. 4.**
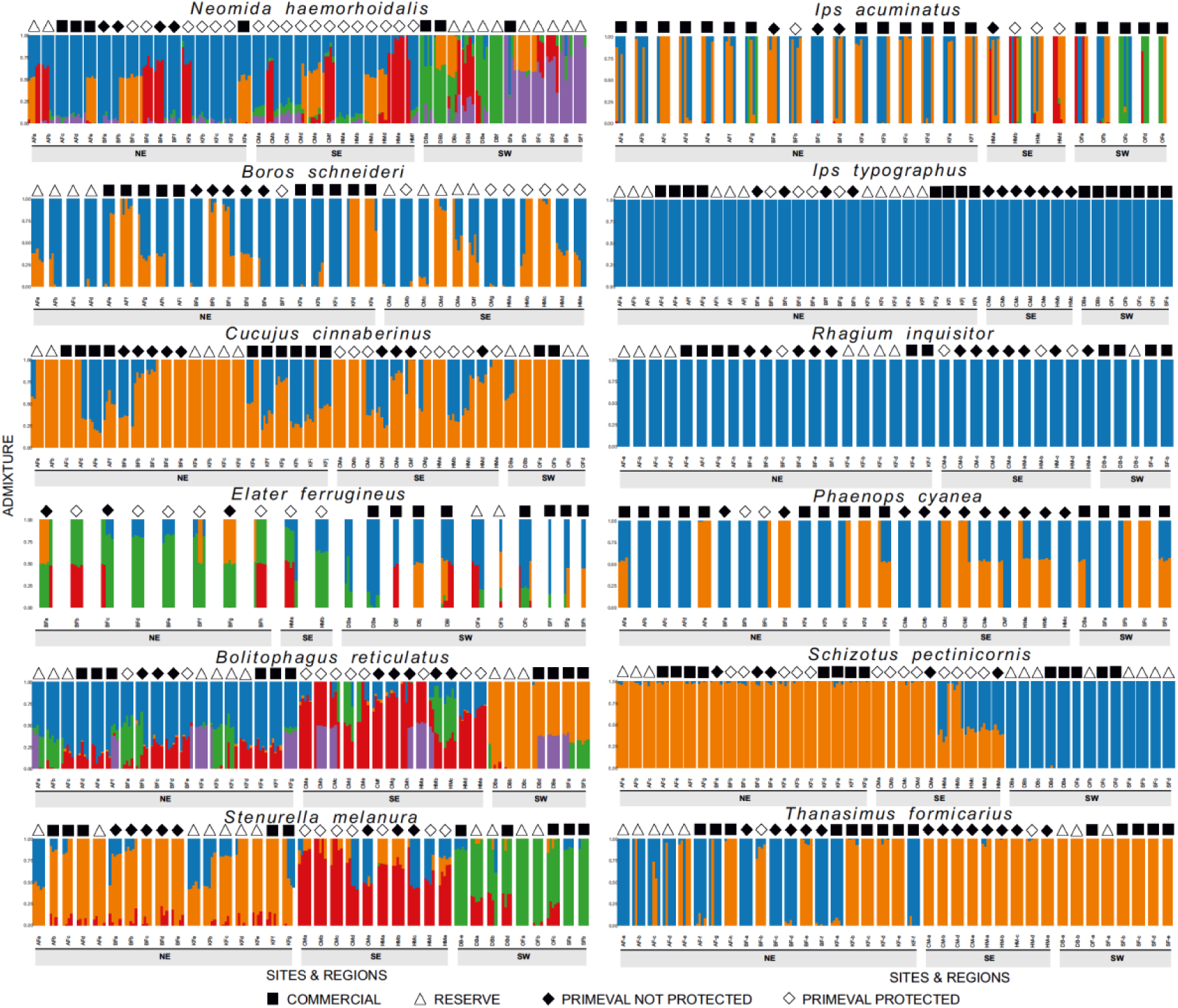
Genetic structure represented by ADMIXTURE based on forests’ geographic regions (NE: north east, SE: south east, SW: south west) and four management levels (commercial, nature reserve, primeval not protected, and primeval protected).

### Genetic structure- Admixture-management

Ancestry estimates are consistent with DAPC proportions. The common species (*B. reticulatus and S. pectinnicornis*), the pest (*I. acuminatus*), and the rare (*N. hamorrhoidalis*) showed the greatest structure with coherent ancestry blocks with limited cross-assignments among management levels, especially primeval protected and commercial (Fig. 4). The rare species (*E. ferrugineus*) showed moderate genetic structure (Fig. 4). In contrast, the remaining common/pest taxa (*T. formicarius*, *R. inquisitor*, *I. typographus*) and the rare taxa (*C. cinnaberinus* and *B. schneideri*) showed more mixed ancestry across management levels (Fig. 4). *pairwise Fst- heatmap-geography*

The pairwise F_st_ heatmap reveals pronounced east-west contrasts in several species, while a few taxa display north-south separation (e.g., *R. inquisitor* and *E. ferrugineus*) (Fig. S2). The east-west pattern was most evident in some common beetles (e.g., *S. pectinicornis, T. formicarius,* and *B. reticulatus*) and in rare species (e.g., *N. haemorrhoidalis* and *E. ferrugineus*) (Fig. S2). The rest of the taxa showed weak variance among geography region populations (Fig. S2).

### pairwise Fst- heatmap-management

In the pairwise F_st_ heatmap, several taxa displayed detectable differences among management levels, with the strongest contrasts typically between primeval protected and commercial stands (Fig. S3). This pattern was most evident in some common beetles (e.g., *S. pectinicornis* and *B. reticulatus*) and in rare species (e.g., *E. ferrugineus* and *C. cinnaberinus*) (Fig. S3). By contrast, conifer taxa (*I. typographus*, *I. acuminatus*) and the conifer-oligophag *R. inquisitor* mostly showed weak among-management variance among levels (Fig. S3).

## Discussion

The major finding of this study is that genetic structure of saproxylic beetle populations is mostly shaped by geography (, and less by forest management types. Majority of species, except some wood generalists and pests, have genetically differentiated populations due to isolation by distance. Rare taxa and common specialist of old-growth forests showed also genetic distinctiveness of their populations in protected than in commercial forests, whereas remaining species, including pests, are not affected by wood logging.

### West-to-east differentiation of populations

The findings of this study indicate that geographic distance explained a strong proportion of genetic variation in many taxa, especially common species. Genetic structure, as shown by DAPC and admixture (by region), revealed strong east-west clustering in six species (three common: *S. pectinicornis*, *B. reticulatus*, and *S. melanura;* one pest: *P. cyanea*; two rare: *C. cinnaberinus* and *N. haemorrhoidalis*), which supports the first hypothesis. This geographic structuring is consistent with classic isolation-by-distance (IBD) patterns (Van Strien et al. 2015; Chang et al. 2022; Hancock et al. 2024) and reflects limited gene flow over long distances. This is in line with previous work on saproxylic beetles, which shows strong spatial genetic structure, especially among niche-specialized and less vagile species (Oleksa et al. 2013; Drag et al. 2015; Eberle et al. 2021). For example, Oleksa et al. (2013) found that two tree-hollow-associated beetles (*E. ferrugineus* and *Osmoderma barnabita*) showed strong kin-based spatial autocorrelation, with the specialist species revealing more pronounced structure than the generalists. Although Oleksa et al. (2013) examined spatial genetics on a local scale, this pattern parallels what we see in our results, where common taxa still show significant geographic structure, but the effect is stronger in those common taxa rather than weak or absent in rare species (except *N. haemorrhoidalis*, which showed moderate R^2^=0.228). Moreover, reviews of saproxylic beetle genetics emphasize that limited dispersal and patchy deadwood habitats favour IBD, especially in older, stable forests (Komonen and Müller 2018; Haeler et al. 2021; Kajtoch et al. 2022). For example, across Europe, *Rosalia alpina* exhibited geographically coherent clades and a significant IBD pattern consistent with the restricted dispersal and dependence on older forest deadwood during and after post-glacial dynamics (Drag et al. 2015). Moreover, populations of *Osmoderma eremita*, specialists on old hollow oaks, show limited inter-tree movement and metapopulation dynamics over decades, conditions that promote fine-scale genetic structure and isolation-by-distance in a stable substrate (Ranius 2000). While most species exhibited east-west isolation, *R. inquisitor* and *E. ferrugineus* showed a different pattern, with north-south clustering and partial mixing across latitude. This anisotropic structure suggests that gene flow is directionally constrained, likely by a combination of ecological and historical processes that align with latitude rather than longitude. Several mechanisms could explain the emergence of this (north-south) genetic pattern. First, a post-glacial recolonization legacy can plausibly generate north-south isolation if lineages expanded along distinct latitudinal routes after the last glacial maximum, then met in secondary contact zones (Hewitt 1999; Drag et al. 2015; Cox et al. 2019). Such histories can leave shallow but persistent genomic clusters that run north-south even when contemporary dispersal is not overtly limited. Second, isolation by environment where latitude covaries with season length, temperature, and humidity, which shape wood-decay dynamics, host-tree composition, and pheromone phenology; key factors for saproxylic beetles (Oleksa et al. 2013; Kajtoch et al. 2022). For *R. inquisitor*, a conifer-associated longhorn, latitudinal shifts in conifer dominance and stand age (and thus decay stage of deadwood) could similarly structure populations along latitude more than longitude (Victorsson 2012; Skrzecz et al. 2016; Giannetti et al. 2018). For *E. ferrugineus*, which depends on large veteran broadleaves and hollow-tree microhabitats, well-documented latitudinal gradients in veteran tree continuity can restrict north-south connectivity even across modest geographic distances (Musa et al. 2013; Andersson et al. 2014), with moderate management-driven structure between primeval protected and managed habitats, suggesting additive effects (latitude-linked habitat quality/continuity and local stand history). It is important to note that limited IBD in some taxa, including species, appears to contradict some previous studies, e.g., on *C. cinnaberinus* (Sikora et al. 2023), although the geographic scale differs. Phylogeographic studies over a wide range of saproxylic beetle species often support IBD, whereas this study focused on only a limited part of their ranges in Central Europe. Consequently, isolation could hardly be detected solely on the basis of geographic distance.

Overall, our results confirm that geographic distances and spatial connectivity are the primary forces shaping population genetic differentiation, with limited but detectable anisotropies in certain taxa. The observed east-west and north-south genetic clustering indicates peripheral populations represent distinct genetic units shaped by both geographic distance and historical landscape processes. Central European forests could act as a transient zone in the genetic structuring of saproxylic beetle populations.

### The forest status impact on genetics

Our dbRDA analysis revealed that the unique management contribution (management effect corrected for geography) was generally low, and mostly non-significant across species except for a weak trend in *R. inquisitor* and *P. cyanea* (p ∼ 0.09). This suggests that management explains only a small portion of the variance in genetic differentiation. However, the distance-decay analyses (MMRR), DAPC, and admixture revealed significant management effects in some species. Thus, the overall second hypothesis receives partial support, with a subset of species (particularly rare and common old-growth forest specialist) showing a clear genetic structure across management levels (especially between primeval protected and commercial). Yet in many taxa (notably the common and pests being either generalists or adapted to tree species favored in commercial forests), management differences don’t translate into strong genetic structuring.

This pattern emphasizes that even widespread species can exhibit management-driven structure when they are ecologically specialized on microhabitats disproportionately represented in primeval habitats, such as late-decay substrates or large-diameter deadwood (Seibold et al. 2016; Komonen and Müller 2018). Although these species occur broadly, they often depend on old-growth or continuity-dependent resources, such as hollow trees and coarse woody debris, which are scarce in intensively managed forests (Stokland et al. 2012; Parisi et al. 2018). Consequently, their effective dispersal may be limited by habitat connectivity rather than geographic range. Similar patterns have been documented for other nominally widespread saproxylic beetles showing fine-scale genetic differentiation linked to habitat continuity (Oleksa et al. 2013; Seibold et al. 2016; Kajtoch et al. 2022).

Therefore, the observed management effect in these common species is not paradoxical but reflects ecological specialization within widespread distributions, a scenario where general geographic ubiquity masks dependence on rare microhabitats (Seibold et al. 2016). In contrast, the more disturbance-tolerant conifer taxa (*R. inquisitor* (conifer-generalist), *I. typographus* (spruce specialist), *I. acuminatus,* and *P. cyanea* (pine specialists)) showed wide overlap across management regimes, consistent with their ability to exploit younger or disturbed habitats and disperse effectively across landscapes (Victorsson 2012; Netherer et al. 2021; Papek et al. 2024). Thus, our results align with ecological expectations once differences in habitat specialization and effective dispersal are considered. This generates a question of why the management effect might be muted in some taxa?

First, if dispersal is high, gene flow may quickly homogenize populations across management regimes. For example, our pest *I. typographus* probably has high connectivity across forest landscapes (Netherer et al. 2021), and also rare *C. cinnaberinus* expresses low genetic differentiation across its range in Europe (Sikora et al. 2023), likely due to recent expansion (Horák et al. 2012). Second, if commercial and primeval forests (as well as protected and not-protected stands), are relatively close (in spatial context) and not completely isolated, landscape connectivity can maintain gene flow, then management may act more as a fine-scale environmental filter rather than a genetic barrier, consistent with landscape genetic theory that permeability of nearby habitats mosaics often outweighs categorical habitats labels (Storfer et al. 2010; Lowe et al. 2015; Winiger et al. 2023). Third, the time scale of differentiation: genetic structure occurs gradually, particularly in long-lived organisms or those of overlapping generations; if management changes are recent relative to generation times, the genetic signal may lag (Landguth et al. 2010; Lowe et al. 2015). Fourth, where deadwood continuity and structure complexity are maintained even in commercial forests, the genetic consequences may be reduced (Seibold et al. 2015, 2016). Fifth, the sampling design for this study assumed collecting beetles from four discrete forest types, whereas in nature, both environmental conditions (e.g., deadwood availability) and population genetic structure are continuous. Therefore, detection of significant relations between forest quality (naturalness) and beetle genetic diversity would require different strategy for analyses (ongoing study, unpublished). Beetle sampling in this study was intended to collect species also in typical commercial forests, although in reality it turned out to be challenging, as even common (incl. pests) could not be found in stands with no deadwood resources (which is typical for Central European commercial forests; Vítková et al. 2018). Conversely, some species, particularly conifer-specialized pests, were hard to collect in protected stands, as there are mostly broadleaved trees. Reviews for saproxylic organisms emphasize that the qualitative nature of deadwood habitats (e.g., connectivity, size, decay level) matters greatly (Stokland et al. 2012; Parisi et al. 2018). If commercial forests retain sufficient structural complexity, genetic isolation may be limited.

In rare species, the moderate management-driven structure suggests that these taxa may be more sensitive to forest history differences. For all selected rare beetle taxa (*B. schneideri, C. cinnaberinus, E. ferrugineus, N. haemorrhoidalis*), some analyses (particularly DAPC plots) showed the strongest distinctiveness of beetles collected in primeval stands and in remaining forests (commercial and nature reserves localized within commercial forests). Moreover, in all these species, populations from protected and not-protected primeval forests are at least partially differentiated. These findings are in line with expectations that populations of rare (threatened, relics) are strongly affected by forest naturalness (including history of management or conservation). This applied also to some still common species being specialists of old-growth forests (e.g. *S. pectinicornis*, *B. reticulatus*). Although small sample sizes, lower connectivity (limited gene flow), and potentially recent declines (caused by past forest management and deficiency of deadwood in forests prior to protection), weaken the signal (Andersson et al. 2014; Bełcik et al. 2019)

In our results, the strongest management-driven structure appeared in those species that already showed strong geographic structuring (common and rare). This suggests that management effects may amplify existing geographic structure by further limiting connectivity in certain forest types. For example, in *S. pectinicornis* and *B. reticulatus*, primeval protected sites may act as refugia with reduced disturbance, higher and continuous deadwood supply, and hence potentially larger effective population size; in commercial forests, fragmentation of coarse woody debris and greater isolation may reduce effective gene flow and raise drift (Jonsson et al. 2003; Parisi et al. 2018; Oettel et al. 2023).

Thus, the fact that management’s broader effects appear modest reinforces the broader theme that for many forest-dependent invertebrates, geography and dispersal limitation dominate structuring, with management acting as a secondary driver. This echoes findings in other taxa (e.g., the resilience of forest fragmentation genetics (Lowe et al., 2015), which show that structural connectivity and gene-flow can partially buffer management/fragmentation effects. In sum, while there is clear evidence of management-driven genetic structure in some species, the effect is subtle and context dependent. These issues are goals of ongoing study.

### Implications on conservation and management

Indeed, our results provide actionable conservation insights. The clear geographic structure in many species means that populations at either extreme (east-west or north-south) may be effectively genetically isolated; for example, structuring in some common species (*S. pectinicornis*, *B. reticulatus*, and *S. melanura*), as well as in some rare (*E. ferrugineus*, *N. haemorrhoidalis*), suggested populations separated by regions may represent distinct management units (MUs) or even evolutionary significant units (ESUs) (Knutsen et al. 1999; Funk et al. 2012; Zamoroka et al. 2022). The same cannot be concluded for *C. cinnaberinus* likely due to its recent recovery (Horák et al. 2012; Sikora et al. 2023), or for rare *B. schneideri*, which is present only in eastern Poland. The management-driven structure in some species highlights that populations in commercial forests (especially those adjacent to or within primeval protected forests) may be genetically distinct and potentially subject to reduced gene flow or effective population size. From a practical standpoint, populations in primeval protected forests often maintain higher connectivity and continuity of deadwood habitats, and they may serve as genetic sources (Paillet et al. 2010; Busse et al. 2022; Haeler et al. 2024). In contrast, populations in commercial forests may be more isolated or subject to drift, as suggested by our results for species showing management-driven structure (Aravanopoulos 2018; Schlaepfer et al. 2018; Cox et al. 2020) and are likely to act as sinks (Pulliam 1988). This suggests that conservation actions should prioritise: 1- maintaining or enhancing connectivity between populations (especially across regions), 2- protecting primeval or restoring close-to-natural (old-growth) forest stands (particularly those currently isolated in managed landscapes), and 3- focusing on those species showing both strong genetic structure and management sensitivity (rare-relics, also some specialized common taxa) as high-priority. Recent reviews emphasise the importance of such targeted genetic monitoring for saproxylic species, particularly given their habitat specialisation and susceptibility to fragmentation (Kajtoch et al. 2022; Krovi et al. 2025).

Moreover, distance-decay results emphasizing increased genetic differentiation at shorter distances for certain management levels suggest that genetic isolation may already be occurring at relatively fine spatial scales (within 10s - 100s km). Thus, even seemingly proximate populations may require corridors or stepping-stone connectivity (e.g., by retention of deadwood resources, retention of old-growth elements within commercial forests). This aligns with ecological theory emphasising that deadwood-dependent and old-growth habitat specialist organisms often display restricted dispersal and strong fine-scale structuring (Schmuki et al. 2006; Kajtoch et al. 2022).

Finally, our findings open the possibility of management units that integrate both geography and forest management. For instance, populations in a given region (for example, NE) living in commercial forests may differ genetically from those in the same region but inhabiting a primeval protected forest. These could thus represent distinct units for monitoring and management. The fact that admixture and DAPC reveal distinct ancestry blocks for some species by region and/or management means there could be cryptic genetic subdivisions that warrant recognition in conservation planning. Given the often-overlooked genetic component in saproxylic beetle conservation, our dataset provides concrete genetic evidence to inform forest management policy (e.g., selection of priority stands, identification of gene-flow bottlenecks, and design of dead-wood retention strategies).

### Summary and conclusion

In summary, our multi-species genomic study of 12 saproxylic beetles demonstrates that geography (geographic distance among populations) is the primary driver of genetic differentiation, with forest management regime playing a secondary but detectable role in some species. The clear east-west regional clustering among some species supports our first hypothesis of significant geographic differentiation. The second hypothesis, that populations from primeval vs commercial forests are distinct, is supported for a subset of taxa, though management effects are modest overall. Finally, the dataset does indeed support conservation applications: by revealing populations with strong genetic isolation or distinct ancestry blocks (by region or management), it provides a basis for prioritising connectivity conservation and the protection of high-value forest stands. These patterns are summarized in Fig. 5.

**Fig. 5.**
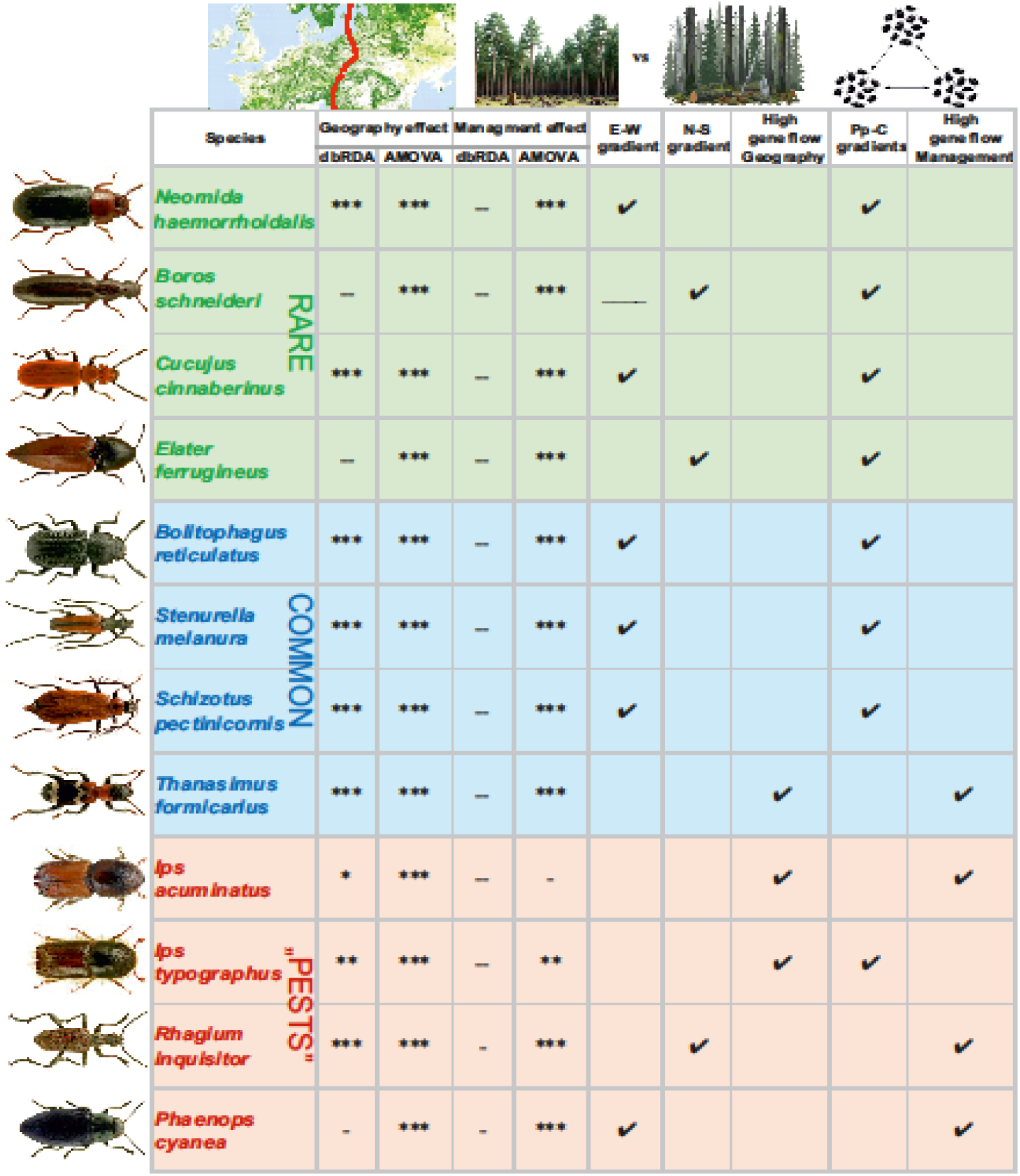
Summary of the effects of geography, management, spatial and management gradients, and gene flow patterns detected in saproxylic beetles. Species groups are colour-coded as follows: common (blue), pests (red), and rare (green). Significance levels are indicated by symbols: *** p< 0.0001, ** p<0.001, * p<0.01, - p <0.1 (weak trend), -- not significant, and _____ populations from west not present. Patterns of spatial and management gradients are abbreviated as E-W: east-west, N-S: north-south, and Pp-M: primeval protected-commercial

From a practical standpoint, the results advise that conservation of saproxylic beetle genetic diversity should emphasise regional connectivity (i.e., ensure gene flow across the regions) and forest management that retains continuous large deadwood substrates and avoids isolation of stands. For less mobile, more habitat-specialised species, the distinction between primeval protected vs commercial forest is meaningful: prioritising primeval protected stands as refugia and linking managed stands to these by retaining structural complexity may be especially beneficial. At the same time, for more mobile or generalist taxa, the management regime appears less critical, and connectivity at the landscape scale likely remains functional.

Overall, our study adds to the growing body of literature documenting the importance of spatial and ecological factors in shaping the genetic structure of saproxylic beetles (Komonen and Müller 2018; Kajtoch et al. 2022) and emphasises that while forest management matters, the spatial scale of dispersal and gene flow is often the dominant factor. By combining high-resolution genomic data with both geographic and management stratification, we provide actionable insights into the conservation genetics of saproxylic beetles and underscore the value of integrating genetic monitoring into forest biodiversity management.

## Acknowledgements

This study was financed by a grant from the National Science Center Poland (UMO-2021/43/B/NZ9/00991, PI – Łukasz Kajtoch).

## Ethical statement

Collection of protected beetle species and in protected areas was executed according to permissions granted by Polish Ministry of Environment (DOP-1.61.61.2021.TP 1516071.4994962.3977511 and DOP-1.61.61.2021.TP 1516071.4994586.3977515), General Directorate of Environmental Protection (DZP-WF.6401.68.2021.AS) and Regional Directorates of Environmental Protection in Białystok (WPN.6205.74.2021.MM), Rzeszów (WPN.6205.116.2021.MP.2 and WPN.6205.114.2021.ŁL.2), Kielce (WPN.I.6205.1.49.2021.EJP.2) and Wrocław (WPN.6205.143.2021.AR).

## References

Abdullah SA, Nakagoshi N (2007) Forest fragmentation and its correlation to human land use change in the state of Selangor, peninsular Malaysia. For Ecol Manage 241:39–48. 10.1016/j.foreco.2006.12.016

Aguirre NC, Filippi CV, Zaina G, et al (2019) Optimizing DDRADseq in non-model species: A Case Study in Eucalyptus dunnii Maiden. Agronomy 9:484. 10.3390/agronomy9090484

Alexander DH, Novembre J, Lange K (2009) Fast model-based estimation of ancestry in unrelated individuals. Genome Res 19:1655–1664. 10.1101/gr.094052.109

Andersson K, Bergman KO, Andersson F, et al (2014) High-accuracy sampling of saproxylic diversity indicators at regional scales with pheromones: The case of Elater ferrugineus (Coleoptera, Elateridae). Biol Conserv 171:156–166. 10.1016/j.biocon.2014.01.007

Aravanopoulos FA (2018) Do silviculture and forest management affect the genetic diversity and structure of long-impacted forest tree populations? Forests 9:355.

Audisio P, Brustel H, Carpaneto GM, et al (2009) Data on molecular taxonomy and genetic diversification of the European Hermit beetles, a species complex of endangered insects (Coleoptera: Scarabaeidae, Cetoniinae, Osmoderma). Journal of Zoological Systematics and Evolutionary Research 47:88–95. 10.1111/j.1439-0469.2008.00475.x

Barbier EB, Burgess JC, Grainger A (2010) The forest transition: Towards a more comprehensive theoretical framework. Land use policy 27:98–107. 10.1016/j.landusepol.2009.02.001

Barbosa S, Mestre F, White TA, et al (2018) Integrative approaches to guide conservation decisions: Using genomics to define conservation units and functional corridors. Mol Ecol 27:3452–3465. 10.1111/mec.14806

Bayat M, Bettinger P, Heidari S, et al (2021) A combination of biotic and abiotic factors and diversity determine productivity in natural deciduous forests. Forests 12:1450. 10.3390/f12111450

Bayona-Vásquez NJ, Glenn TC, Kieran TJ, et al (2019) Adapterama III: Quadruple-indexed, double/triple-enzyme RADseq libraries (2RAD/3RAD). PeerJ 7:e7723. 10.7717/peerj.7724

Bełcik M, Goczał J, Ciach M (2019) Large-scale habitat model reveals a key role of large trees and protected areas in the metapopulation survival of the saproxylic specialist Cucujus cinnaberinus. Biodivers Conserv 28:3851–3871. 10.1007/s10531-019-01854-0

Bohn FJ, Huth A (2017) The importance of forest structure to biodiversity-productivity relationships. R Soc Open Sci 4: 160521. 10.1098/rsos.160521

Burns M, Starrett J, Derkarabetian S, et al (2017) Comparative performance of double-digest RAD sequencing across divergent arachnid lineages. Mol Ecol Resour 17:418–430. 10.1111/1755-0998.12575

Busse A, Cizek L, Čížková P, et al (2022) Forest dieback in a protected area triggers the return of the primeval forest specialist Peltis grossa (Coleoptera, Trogossitidae). Conserv Sci Pract 4:612. 10.1111/csp2.612

Casacci LP, Barbero F, Balletto E (2014) The “Evolutionarily Significant Unit” concept and its applicability in biological conservation. Italian Journal of Zoology 81:182–193. 10.1080/11250003.2013.870240

Chang CW, Fridman E, Mascher M, et al (2022) Physical geography, isolation by distance and environmental variables shape genomic variation of wild barley (Hordeum vulgare L. ssp. spontaneum) in the Southern Levant. Heredity (Edinb) 128:107–119. 10.1038/s41437-021-00494-x

Chave J, Muller-Landau HC, Baker TR, et al (2006) Regional and phylogenetic variation of wood density across 2456 neotropical tree species. Ecological Applications 16:2356–2367. 10.1890/1051-0761(2006)016[2356:RAPVOW]2.0.CO;2

Christensen M, Emborg J (1996) Biodiversity in natural versus managed forest in Denmark

Coates BS, Bayles DO, Wanner KW, et al (2011) The application and performance of single nucleotide polymorphism markers for population genetic analyses of lepidoptera. Front Genet 2:38. 10.3389/fgene.2011.00038

Cox K, McKeown N, Antonini G, et al (2019) Phylogeographic structure and ecological niche modelling reveal signals of isolation and postglacial colonisation in the European stag beetle. PLoS One 14:e215860. 10.1371/journal.pone.0215860

Cox K, McKeown N, Vanden Broeck A, et al (2020) Genetic structure of recently fragmented suburban populations of European stag beetle. Ecol Evol 10:12290–12306. 10.1002/ece3.6858

de Groot M, Diaci J, Ogris N (2019) Forest management history is an important factor in bark beetle outbreaks: Lessons for the future. For Ecol Manage 433:467–474. 10.1016/j.foreco.2018.11.025

De Vivo M, Chou MH, Wu SP, et al (2023) Genomic tools for comparative conservation genetics among three recently diverged stag beetles (Lucanus, Lucanidae). Insect Conserv Divers 16:853–869. 10.1111/icad.12678

Drag L, Hauck D, Bérces S, et al (2015) Genetic differentiation of populations of the threatened saproxylic beetle Rosalia longicorn, Rosalia alpina (Coleoptera: Cerambycidae) in Central and South-east Europe. Biological Journal of the Linnean Society 116:911–925. 10.1111/bij.12624

Eberle J, Husemann M, Doerfler I, et al (2021) Molecular biogeography of the fungus-dwelling saproxylic beetle Bolitophagus reticulatus indicates rapid expansion from glacial refugia. Biological Journal of the Linnean Society, 2021, 133 (3), pp.766–778

Ellerstrand SJ, Choudhury S, Svensson K, et al (2022) Weak population genetic structure in Eurasian spruce bark beetle over large regional scales in Sweden. Ecol Evol 12:e9078. 10.1002/ece3.9078

Excoffier L, Smouse PE, Quattro JM (1992) Analysis of Molecular Variance Inferred From Metric Distances Among DNA Haplotypes: Application to Human Mitochondrial DNA Restriction Data. Genetics 131: 479–491

Fahrig L (2003) Effects of Habitat Fragmentation on Biodiversity. Source: Annual Review of Ecology, Evolution, and Systematics 34:487–515. 10.1146/132419

Feng M, Sexton JO, Huang C, et al (2016) Earth science data records of global forest cover and change: Assessment of accuracy in 1990, 2000, and 2005 epochs. Remote Sens Environ 184:73–85. 10.1016/j.rse.2016.06.012

Fraser DJ, Bernatchez L (2001) Adaptive evolutionary conservation: towards a unified concept for defining conservation units. Mol Ecol 10:2741–2752. 10.1046/j.0962-1083.2001.01411.x

Freer-Smith P, Muys B, Bozzano M, et al (2019) Plantation forests in Europe: challenges and opportunities. From Science to Policy 9.European Forest Institute. 10.36333/fs09

Funk WC, McKay JK, Hohenlohe PA, Allendorf FW (2012) Harnessing genomics for delineating conservation units. Trends Ecol Evol 27:489–496

Giannetti F, Barbati A, Mancini LD, et al (2018) European Forest Types: toward an automated classification. Ann For Sci 75:6. 10.1007/s13595-017-0674-6

Giesecke T, Brewer S (2018) Notes on the postglacial spread of abundant European tree taxa. Veg Hist Archaeobot 27:337–349. 10.1007/s00334-017-0640-0

Gillespie TW, Lipkin B, Sullivan L, et al (2012) The rarest and least protected forests in biodiversity hotspots. Biodivers Conserv 21:3597–3611. 10.1007/s10531-012-0384-1

Gimmel ML, Ferro ML (2018) General Overview of Saproxylic Coleoptera. pp 51–128

Goslee SC, Urban DL (2007) The ecodist package for dissimilarity-based analysis of ecological data. J Stat Softw 22:1–19. 10.18637/jss.v022.i07

Grove SJ (2002a) Saproxylic insect ecology and the sustainable management of forests. Annu Rev Ecol Syst 33:1–23

Grove SJ (2002b) Saproxylic insect ecology and the sustainable management of forests. Annu Rev Ecol Syst 33:1–23

Guo SJ, Chai D-D, Guo S-J, et al (2013) The Major Factors Affecting Ectomycorrhizal Fungi Diversity in the Forest Ecosystem. Advance Journal of Food Science and Technology 5:879–890

Haeler E, Bergamini A, Blaser S, et al (2021) Saproxylic species are linked to the amount and isolation of dead wood across spatial scales in a beech forest. Landsc Ecol 36:89–104. 10.1007/s10980-020-01115-4

Haeler E, Stillhard J, Hindenlang Clerc K, et al (2024) Dead wood distributed in different-sized habitat patches enhances diversity of saproxylic beetles in a landscape experiment. Journal of Applied Ecology 61:316–327. 10.1111/1365-2664.14554

Hajibabaei M, Singer GAC, Hebert PDN, Hickey DA (2007) DNA barcoding: how it complements taxonomy, molecular phylogenetics and population genetics. Trends in Genetics 23:167–172. 10.1016/j.tig.2007.02.001

Hancock ZB, Toczydlowski RH, Bradburd GS (2024) A spatial approach to jointly estimate Wright’s neighborhood size and long-term effective population size. Genetics 227: iyae094. 10.1093/genetics/iyae094

Hebert PDN, Cywinska A, Ball SL, DeWaard JR (2003) Biological identifications through DNA barcodes. Proceedings of the Royal Society B: Biological Sciences 270:313–321. 10.1098/rspb.2002.2218

Hewitt G (2000) The genetic legacy of the Quaternary ice ages. School of Biological Sciences, University of East Anglia, Norwich NR4 7TJ, UK

Hewitt GM (1999) Post-glacial re-colonization of European biota. Biological Journal of the Linnean Society 68:87–112. 10.1111/j.1095-8312.1999.tb01160.x

Hijmans, R.J. (2023). geosphere: Spherical Trigonometry. R package. 10.1007/s00190012

Hjältén J, Stenbacka F, Pettersson RB, et al (2012) Micro and macro-habitat associations in saproxylic beetles: Implications for biodiversity management. PLoS One 7:. 10.1371/journal.pone.0041100

Horák, J., Chumanová, E., & Hilszczański, J. (2012). Saproxylic beetle thrives on the openness in management: A case study on the ecological requirements of Cucujus cinnaberinus from Central Europe. Insect Conservation and Diversity, 5(6), 403–413.

Jombart T (2008) Adegenet: A R package for the multivariate analysis of genetic markers. Bioinformatics 24:1403–1405. 10.1093/bioinformatics/btn129

Jombart T, Devillard S, Balloux F (2010) Discriminant analysis of principal components: A new method for the analysis of genetically structured populations. BMC Genet 11:94. 10.1186/1471-2156-11-94

Jonsson M, Johannesen J, Seitz A (2003) Comparative genetic structure of the threatened tenebrionid beetle *Oplocephala haemorrhoidalis* and its common relative Bolitophagus reticulatus. Journal of Insect Conservation 7: 111–124.

Kajtoch, Gronowska M, Plewa R, et al (2022) A review of saproxylic beetle intra- and interspecific genetics: current state of the knowledge and perspectives. European Zoological Journal 89:481–501

Kajtoch Ł, Kolasa M, Kubisz D, et al (2019) Using host species traits to understand the Wolbachia infection distribution across terrestrial beetles. Sci Rep 9:847. 10.1038/s41598-018-38155-5

Knutsen H, Arne Rukke Bj, Erik Jorde P, Ims RA (2000) Genetic differentiation among populations of the beetle *Bolitophagus reticulatus* (Coleoptera: Tenebrionidae) in a fragmented and a continuous landscape. Heredity 84:667–676

Komonen A, Müller J (2018) Dispersal ecology of deadwood organisms and connectivity conservation. Conservation Biology 32:535–545

Kozák D, Svitok M, Wiezik M, et al (2021) Historical Disturbances Determine Current Taxonomic, Functional and Phylogenetic Diversity of Saproxylic Beetle Communities in Temperate Primary Forests. Ecosystems 24:37–55. 10.1007/s10021-020-00502-x

Krovi RS, Amer NR, Oczkowicz M, Kajtoch Ł (2024) Meta-analysis of spatial genetic patterns among European saproxylic beetles. Biodiversity and Conservation 34:1–27. 10.1007/s10531-024-02940-8

Kuuluvainen T (2009) Forest management and biodiversity conservation based on natural ecosystem dynamics in Northern Europe: The complexity challenge. Ambio 38:309–315. 10.1579/08-A-490.1

Lagisz M, Port G, Wolff K (2010) A cost-effective, simple and high-throughput method for DNA extraction from insects. Insect Sci 17:465–470. 10.1111/j.1744-7917.2010.01318.x

Landguth EL, Cushman SA, Schwartz MK, et al (2010) Quantifying the lag time to detect barriers in landscape genetics. Mol Ecol 19:4179–4191. 10.1111/j.1365-294X.2010.04808.x

Legendre P, Andersson MJ (1999) Distance-based redundancy analysis: Testing multispecies responses in multifactorial ecological experiments. Ecol Monogr 69:1–24. 10.1890/0012-9615(1999)069[0001:DBRATM]2.0.CO;2

Lowe AJ, Cavers S, Boshier D, et al (2015) The resilience of forest fragmentation genetics--no longer a paradox--we were just looking in the wrong place. Heredity (Edinb) 115:97–99

Magbanua Z V., Hsu CY, Pechanova O, et al (2023) Innovations in double digest restriction-site associated DNA sequencing (ddRAD-Seq) method for more efficient SNP identification. Anal Biochem 662: 115001. 10.1016/j.ab.2022.115001

Mazur A, Witkowski R, Kuźmiński R, et al (2021) The structure of saproxylic beetle assemblages in view of coarse woody debris resources in pine stands of western poland. Forests 12:1558. 10.3390/f12111558

Mori AS, Lertzman KP, Gustafsson L (2017) Biodiversity and ecosystem services in forest ecosystems: a research agenda for applied forest ecology. Journal of Applied Ecology 54:12–27. 10.1111/1365-2664.12669

Murray DL, Peers MJL, Majchrzak YN, et al (2017) Continental divide: Predicting climatemediated fragmentation and biodiversity loss in the boreal forest. PLoS One 12(5): e0176706. 10.1371/journal.pone.0176706

Musa N, Andersson K, Burman J, et al (2013) Using Sex Pheromone and a Multi-Scale Approach to Predict the Distribution of a Rare Saproxylic Beetle. PLoS One 8(6): e66149. 10.1371/journal.pone.0066149

Netherer S, Kandasamy D, Jirosová A, et al (2021) Interactions among Norway spruce, the bark beetle Ips typographus and its fungal symbionts in times of drought. J Pest Sci 94:591–614

Oettel J, Zolles A, Gschwantner T, et al (2023) Dynamics of standing deadwood in Austrian forests under varying forest management and climatic conditions. Journal of Applied Ecology 60:696–713. 10.1111/1365-2664.14359

Ogle K, Pathikonda S, Sartor K, et al (2014) A model-based meta-analysis for estimating species-specific wood density and identifying potential sources of variation. Journal of Ecology 102:194–208. 10.1111/1365-2745.12178

Oleksa A, Chybicki IJ, Gawroński R, et al (2013) Isolation by distance in saproxylic beetles may increase with niche specialization. J Insect Conserv 17:219–233. 10.1007/s10841-012-9499-7

Paillet Y, Bergès L, HjÄltén J, et al (2010) Biodiversity differences between managed and unmanaged forests: Meta-analysis of species richness in Europe. Conservation Biology 24:101–112

Palsbøll PJ, Bérubé M, Allendorf FW (2007) Identification of management units using population genetic data. Trends Ecol Evol 22:11–16. 10.1016/j.tree.2006.09.003

Papek E, Ritzer E, Biedermann PHW, et al (2024) The pine bark beetle Ips acuminatus: an ecological perspective on life-history traits promoting outbreaks. J Pest Sci 97:1093–1122

Parisi F, Pioli S, Lombardi F, et al (2018) Linking deadwood traits with saproxylic invertebrates and fungi in European forests – A review. IForest 11:423–436

Pawson SM, Brin A, Brockerhoff EG, et al (2013) Plantation forests, climate change and biodiversity. Biodivers Conserv 22:1203–1227. 10.1007/s10531-013-0458-8

Peres-Neto PR, Legendre P (2010) Estimating and controlling for spatial structure in the study of ecological communities. Global Ecology and Biogeography 19:174–184. 10.1111/j.1466-8238.2009.00506.x

Perry DA. (1994) Forest ecosystems. Johns Hopkins University Press. Baltimore. Maryland 21218-4319

Peterson BK, Weber JN, Kay EH, et al (2012) Double digest RADseq: An inexpensive method for de novo SNP discovery and genotyping in model and non-model species. PLoS One 7:e37135. 10.1371/journal.pone.0037135

Pulliam HR 1988. “Sources, sinks, and population regulation”. The American Naturalist. 132 (5): 652–61.

R Development Core Team. (2013). R: A Language and Environment for Statistical Computing (Version 4.0.3) [Computer software]. The R Foundation for Statistical Computing, Vienna, Austria. http://www.R-project.org.

Ranius T (2000) Minimum viable metapopulation size of a beetle, Osmoderma eremita, living in tree hollows. Anim Conserv 3:37–43. 10.1111/j.1469-1795.2000.tb00085.x

Ranius T (2006) Measuring the dispersal of saproxylic insects: A key characteristic for their conservation. Popul Ecol (2006) 48:177–188. DOI 10.1007/s10144-006-0262-3

Rasche L, Fahse L, Bugmann H (2013) Key factors affecting the future provision of tree-based forest ecosystem goods and services. Clim Change 118:579–593. 10.1007/s10584-012-0664-5

Rochette NC, Rivera-Colón AG, Catchen JM (2019a) Stacks 2: Analytical methods for paired-end sequencing improve RADseq-based population genomics. Mol Ecol 28:4737–4754. 10.1111/mec.15253

Rochette NC, Rivera-Colón AG, Catchen JM (2019b) Stacks 2: Analytical methods for paired-end sequencing improve RADseq-based population genomics. Mol Ecol 28:4737–4754. 10.1111/mec.15253

Rousset F (1997) Genetic Differentiation and Estimation of Gene Flow from FStatistics Under Isolation by Distance. Genetics 145 1219–1228

Schiegg K (2000) Effects of dead wood volume and connectivity on saproxylic insect species diversity. European Forest Institute 2005.

Schlaepfer DR, Braschler B, Rusterholz HP, Baur B (2018) Genetic effects of anthropogenic habitat fragmentation on remnant animal and plant populations: a meta-analysis. Ecosphere 9(10): e02488. 10.1002/ecs2.2488

Schmitt T (2007) Molecular biogeography of Europe: Pleistocene cycles and postglacial trends. Front Zool 4:11 doi:10.1186/1742-9994-4-11

Schmuki C, Vorburger C, Runciman D, et al (2006) When log-dwellers meet loggers: Impacts of forest fragmentation on two endemic log-dwelling beetles in southeastern Australia. Mol Ecol 15:1481–1492. 10.1111/j.1365-294X.2006.02849.x

Seibold S, Bässler C, Brandl R, et al (2016) Microclimate and habitat heterogeneity as the major drivers of beetle diversity in dead wood. Journal of Applied Ecology 53:934–943. 10.1111/1365-2664.12607

Seibold S, Bässler C, Brandl R, et al (2015) Experimental studies of dead-wood biodiversity - A review identifying global gaps in knowledge. Biol Conserv 191:139–149

Selkoe KA, Toonen RJ (2006) Microsatellites for ecologists: A practical guide to using and evaluating microsatellite markers. Ecol Lett 9:615–629

Siitonen J (2001) Forest Management, Coarse Woody Debris and Saproxylic Organisms: Fennoscandian Boreal Forests as an Example. Ecological Bulletins 49: 11–41.

Simberloff D (1999) The role of science in the preservation of forest biodiversity. For. Ecol. Manag. 115:101–111

Singh VV, Naseer A, Mogilicherla K, et al (2024) Understanding bark beetle outbreaks: exploring the impact of changing temperature regimes, droughts, forest structure, and prospects for future forest pest management. Rev Environ Sci Biotechnol 23:257–290

Skrzecz I, Ukalska J, Tumialis D (2016) Effects of Norway Spruce (Picea abies) Stump Debarking on Insect Colonization in the Polish Sudety Mountains. Mt Res Dev 36:203–212. 10.1659/MRD-JOURNAL-D-15-00073.1

Stokland JN., Siitonen Juha, Jonsson B Gunnar (2012) Biodiversity in Dead Wood. Cambridge University Press. UK.

Storfer A, Murphy MA, Spear SF, et al (2010) Landscape genetics: Where are we now? Mol Ecol 19:3496–3514. 10.1111/j.1365-294X.2010.04691.x

Taberlet P, Fumagalli L, Wust-Saucy AG, Cosson JF (1998) Comparative phylogeography and postglacial colonization routes in Europe. Mol Ecol 7:453–464. 10.1046/j.1365-294x.1998.00289.x

Taubert F, Fischer R, Groeneveld J, et al (2018) Global patterns of tropical forest fragmentation. Nature 554:519–522. 10.1038/nature25508

Taylor AR, Chen HYH, VanDamme L (2009) A review of forest succession models and their suitability for forest management planning. Forest Science 55:23–36. 10.1093/forestscience/55.1.23

Thompson Ian, Mackey Brendan, McNulty Steven, Mosseler Alex (2014) Forest resilience, biodiversity, and climate change: a synthesis of the biodiversity, resilience, stabiblity relationship in forest ecosystems. Secretariat of the convention on the biological diversity, Montreal. Technical Series no. 43, 67 pages.

Thorn S, Förster B, Heibl C, et al (2018) Influence of macroclimate and local conservation measures on taxonomic, functional, and phylogenetic diversities of saproxylic beetles and wood-inhabiting fungi. Biodivers Conserv 27:3119–3135. 10.1007/s10531-018-1592-0

Tóth EG, Köbölkuti ZA, Cseke K, et al (2021) A genomic dataset of single-nucleotide polymorphisms generated by ddRAD tag sequencing in Q. petraea (Matt.) Liebl. populations from Central-Eastern Europe and Balkan Peninsula. Ann For Sci 78:43. 10.1007/s13595-021-01051-6

Ulrich B (1995) The history and possible causes of forest decline in central Europe, with particular attention to the German situation. Reviews 3:262–276. 10.2307/envirevi.3.3-4.262

Van Strien MJ, Holderegger R, Van Heck HJ (2015) Isolation-by-distance in landscapes: Considerations for landscape genetics. Heredity (Edinb) 114:27–37. 10.1038/hdy.2014.62

Victorsson J (2012) Semi-field experiments investigating facilitation: Arrival order decides the interrelationship between two saproxylic beetle species. Ecol Entomol 37:395–401. 10.1111/j.1365-2311.2012.01377.x

Vignal A, Milan D, SanCristobal M, Eggen A (2002) A review on SNP and other types of molecular markers and their use in animal genetics. Genetics Selection Evolution 34:275–305

Wade TG, Riitters KH, Wickham JD, Jones KB (2003) Distribution and Causes of Global Forest Fragmentation. Conserv. Ecol. 7(2): 7

Watson JEM, Evans T, Venter O, et al (2018) The exceptional value of intact forest ecosystems. Nat Ecol Evol 2:599–610

Weir BS, Clark C (1984) Estimating F-Statistics for the Analysis of Population Structure. Evolution, 38(6), pp. 1358–1370

Wenne R (2023) Single Nucleotide Polymorphism Markers with Applications in Conservation and Exploitation of Aquatic Natural Populations. Animals 13, 1089.

Williams JGK, Kubelik AR, Livak KJ, et al DNA polymorphisms amplified by arbitrary primers are useful as genetic markers. Nucleic Acids Res.18–22

Winiger N, Hendel AL, Ganz S, et al (2023) Saproxylic beetles respond to habitat variables at different spatial scales depending on variable type and species’ mobility: the need for multi-scale forest structure management. Biodivers Conserv 32:3355–3377. 10.1007/s10531-023-02663-2

Zamoroka AM, Trócoli S, Shparyk VY, Semaniuk D V. (2022) Polyphyly of the genus Stenurella (Coleoptera, Cerambycidae): Consensus of morphological and molecular data. Biosyst Divers 30:119–136. 10.15421/012212

