## Supplementary file for "Geography and forest naturalness as drivers of genetic diversity in saproxylic beetles"

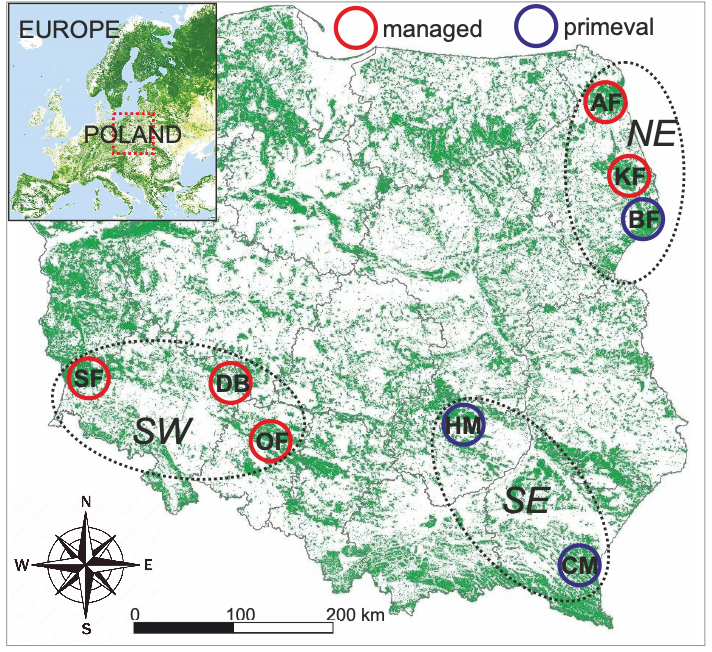

Fig. S1 map of sampling sites of 12 species of saproxylic beetles across Poland.

BF – Białowieża Forest, KF – Knyszyn Forest, AF – Augustów Forest, HM – Holy Cross Mts, CF – Carpathian Mts, OF – Oder Forest, DB – Barych Forest, SF – Silesia Forest.

NE, SE, SW – three regions considered in the study.

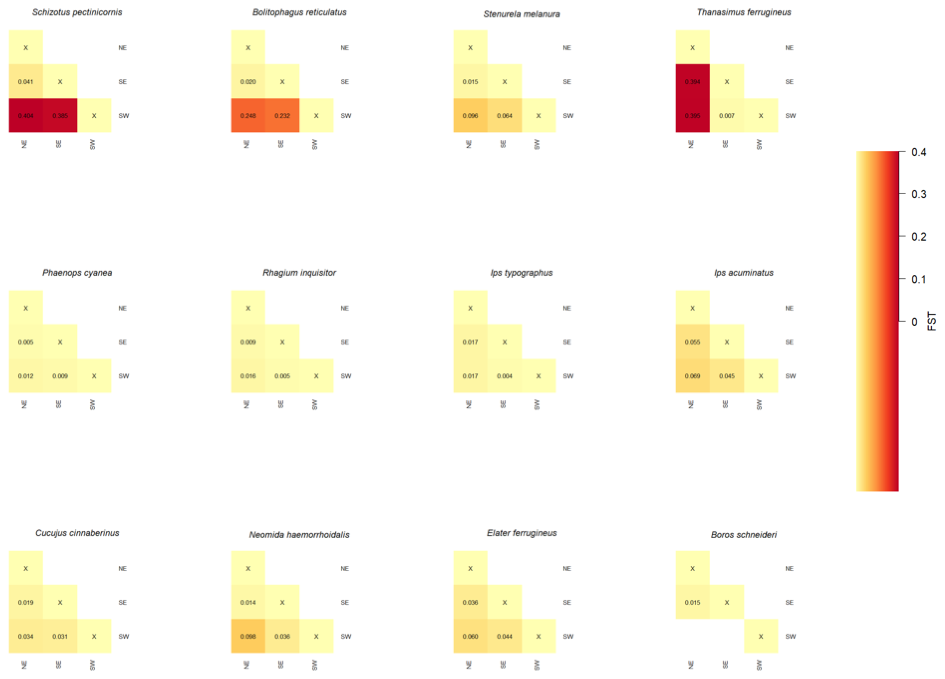

Fig. S2. Genetic structure represented by pairwise Fst heatmap among forests’ geographic regions (NE: north east, SE: south east, SW: south west).

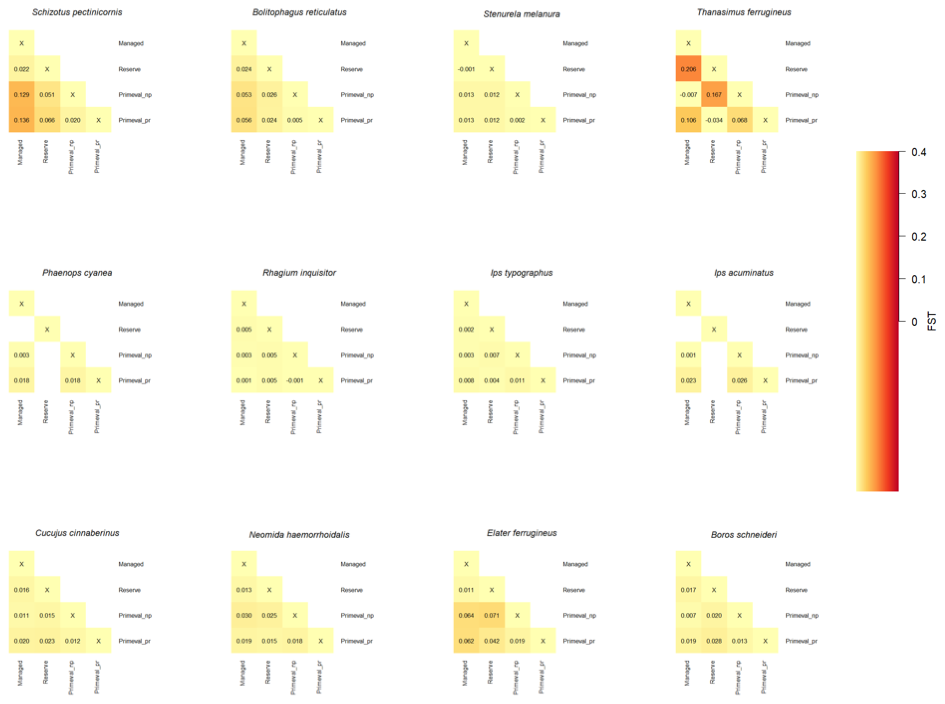

Fig. S3. Genetic structure represented by pairwise Fst heatmap among four forests’ management levels (commercial, nature reserve, primeval not protected, and primeval protected).

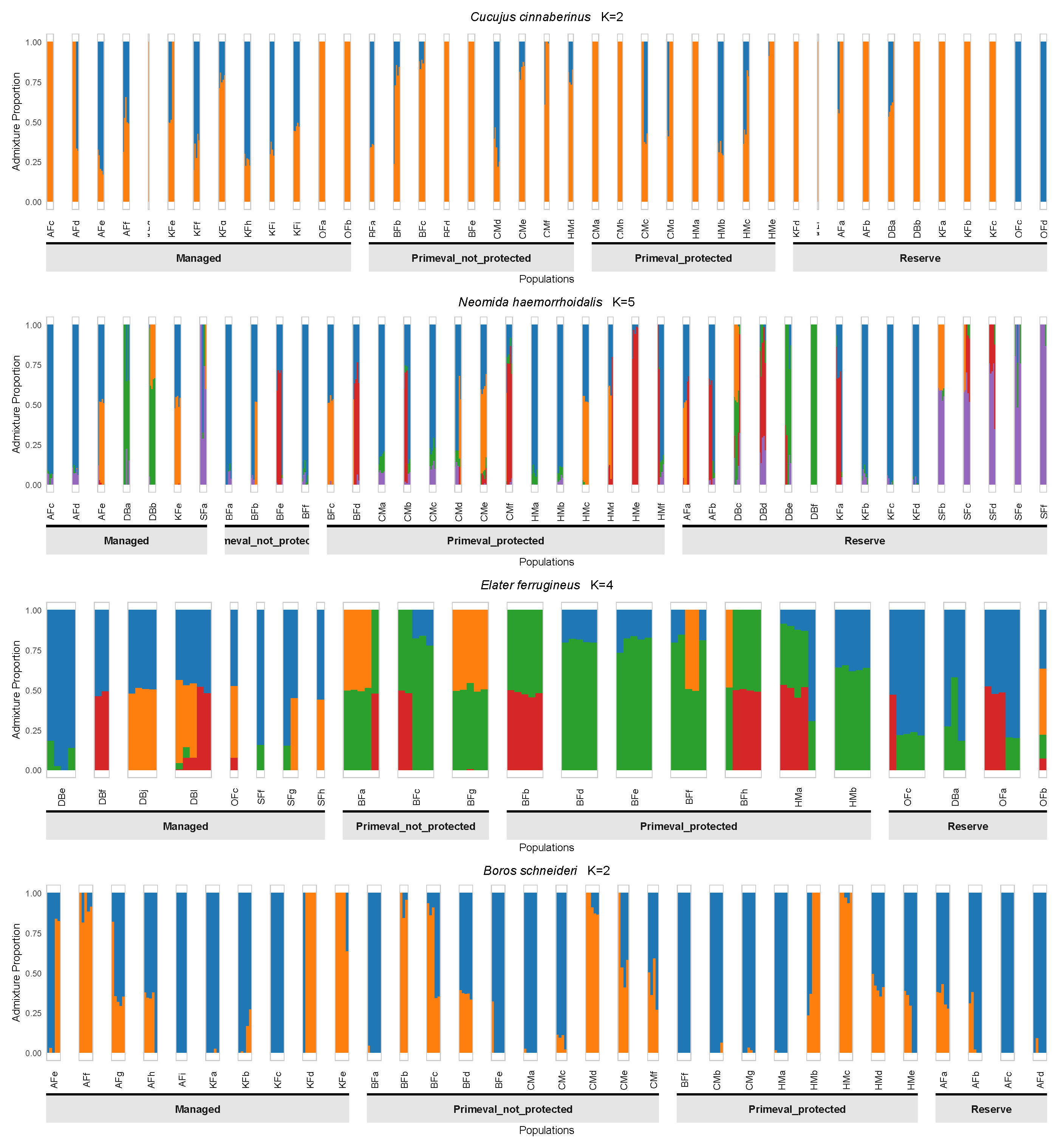

Fig. S4. Genetic structure represented by ADMIXTURE based on four management levels (commercial, nature reserve, primeval not protected, and primeval protected), presented for four beetle species assigned in a category “rare”.

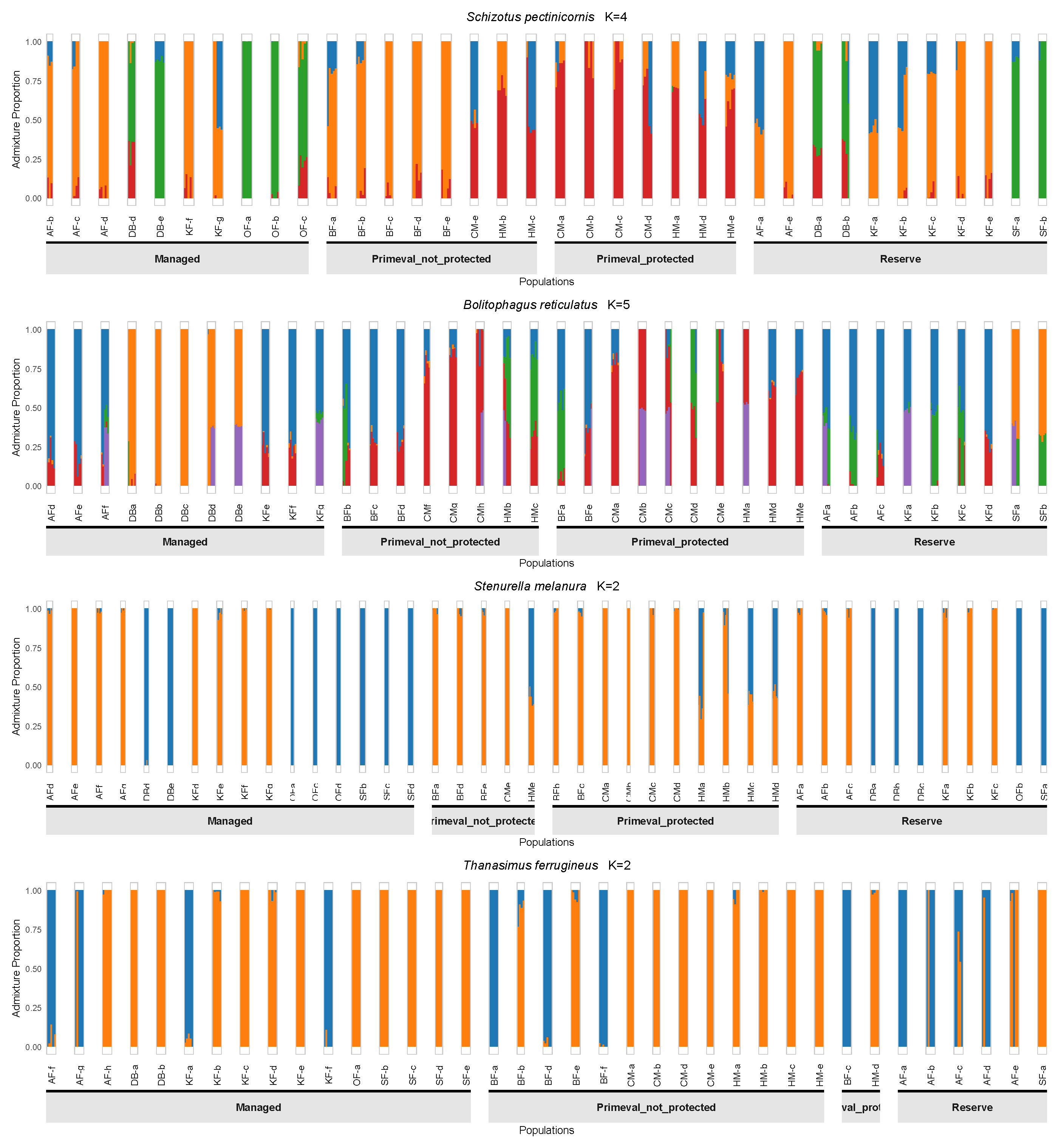

Fig. S5. Genetic structure represented by ADMIXTURE based on four management levels (commercial, nature reserve, primeval not protected, and primeval protected), presented for four beetle species assigned in a category “common”.

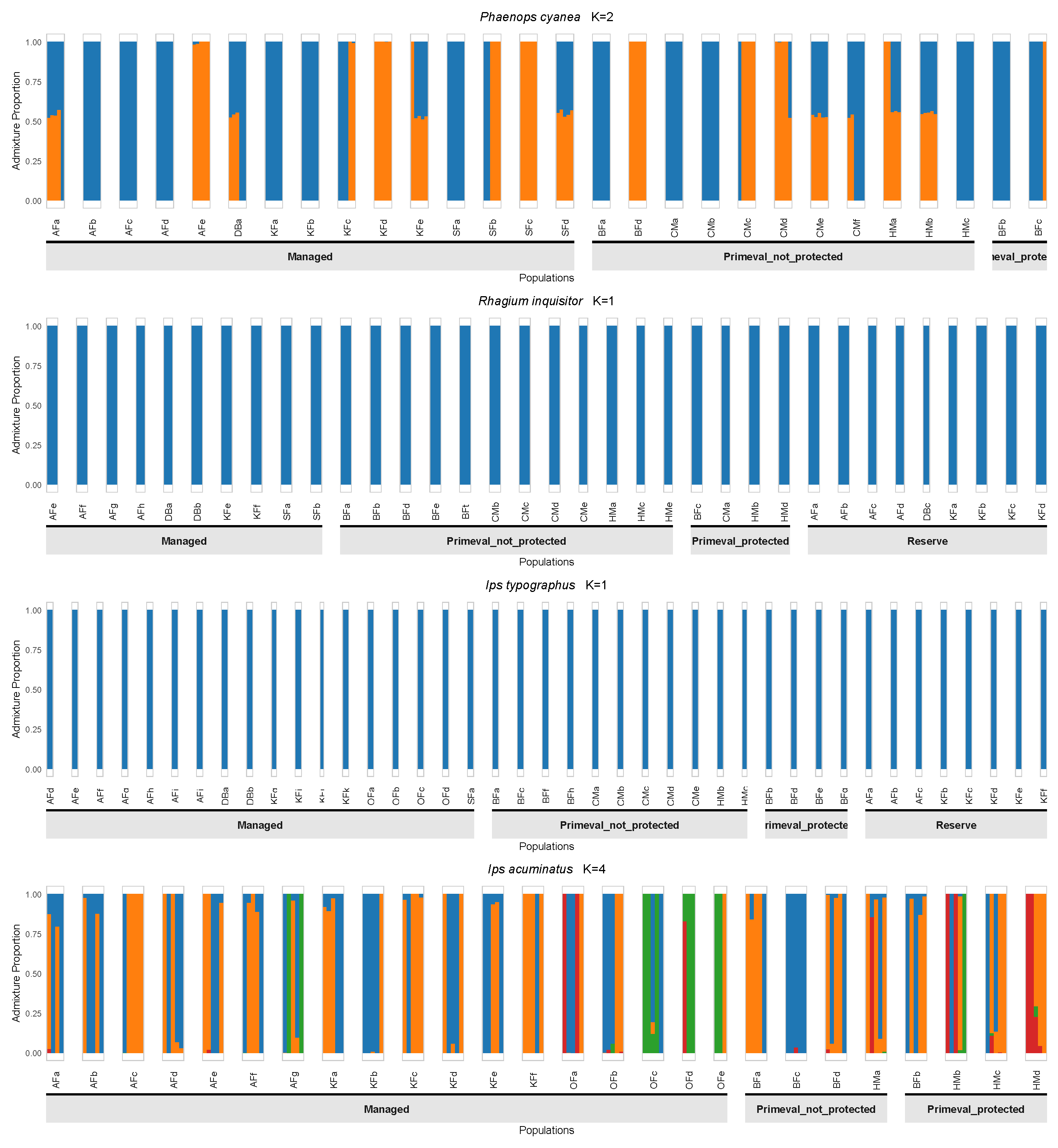

Fig. S6. Genetic structure represented by ADMIXTURE based on four management levels (commercial, nature reserve, primeval not protected, and primeval protected), presented for four beetle species assigned in a category “pest”.

Table S1. Sampling sites of the studied 12 beetle species with corresponding geographic coordinates, forest locations, and management category.

| **Species** | **Forest** | **site** | **Latitude** | **Longitude** | **Geographic region** | **Management** |
| --- | --- | --- | --- | --- | --- | --- |
| *Cucujus cinnaberinus* | BF | CcBFa | 52.686502 | 23.708807 | NE | Primeval_not_protected |
| *Cucujus cinnaberinus* | BF | CcBFb | 52.69084 | 23.727883 | NE | Primeval_not_protected |
| *Cucujus cinnaberinus* | BF | CcBFc | 52.668435 | 23.768888 | NE | Primeval_not_protected |
| *Cucujus cinnaberinus* | BF | CcBFd | 52.687247 | 23.878859 | NE | Primeval_not_protected |
| *Cucujus cinnaberinus* | BF | CcBFe | 52.702839 | 23.889652 | NE | Primeval_not_protected |
| *Cucujus cinnaberinus* | KF | CcKFa | 53.203179 | 23.578818 | NE | Reserve |
| *Cucujus cinnaberinus* | KF | CcKFb | 53.410621 | 23.380008 | NE | Reserve |
| *Cucujus cinnaberinus* | KF | CcKFc | 53.342542 | 23.300179 | NE | Reserve |
| *Cucujus cinnaberinus* | KF | CcKFd | 53.276623 | 23.380327 | NE | Reserve |
| *Cucujus cinnaberinus* | KF | CcKFe | 53.28903 | 23.134141 | NE | Commercial |
| *Cucujus cinnaberinus* | KF | CcKFf | 53.2756 | 23.3047 | NE | Commercial |
| *Cucujus cinnaberinus* | KF | CcKFg | 53.2641 | 23.215724 | NE | Commercial |
| *Cucujus cinnaberinus* | KF | CcKFh | 53.114651 | 23.828271 | NE | Commercial |
| *Cucujus cinnaberinus* | KF | CcKFi | 53.1481 | 23.6824 | NE | Commercial |
| *Cucujus cinnaberinus* | KF | CcKFj | 53.192781 | 23.668165 | NE | Commercial |
| *Cucujus cinnaberinus* | KF | CcKFk | 53.2692 | 23.3188 | NE | Commercial |
| *Cucujus cinnaberinus* | AF | CcAFa | 53.868204 | 23.303332 | NE | Reserve |
| *Cucujus cinnaberinus* | AF | CcAFb | 53.871591 | 23.348753 | NE | Reserve |
| *Cucujus cinnaberinus* | AF | CcAFc | 53.869975 | 23.362405 | NE | Commercial |
| *Cucujus cinnaberinus* | AF | CcAFd | 53.91163 | 23.233723 | NE | Commercial |
| *Cucujus cinnaberinus* | AF | CcAFe | 53.848727 | 23.319015 | NE | Commercial |
| *Cucujus cinnaberinus* | AF | CcAFf | 53.99144 | 23.477051 | NE | Commercial |
| *Cucujus cinnaberinus* | HM | CcHMa | 50.93119 | 20.899131 | SE | Primeval_protected |
| *Cucujus cinnaberinus* | HM | CcHMb | 50.927296 | 20.908708 | SE | Primeval_protected |
| *Cucujus cinnaberinus* | HM | CcHMc | 50.885276 | 21.111485 | SE | Primeval_protected |
| *Cucujus cinnaberinus* | HM | CcHMd | 51.068641 | 20.721844 | SE | Primeval_not_protected |
| *Cucujus cinnaberinus* | HM | CcHMe | 50.965283 | 20.483496 | SE | Primeval_protected |
| *Cucujus cinnaberinus* | CM | CcCMa | 49.695867 | 22.538749 | SE | Primeval_protected |
| *Cucujus cinnaberinus* | CM | CcCMb | 49.656805 | 22.500743 | SE | Primeval_protected |
| *Cucujus cinnaberinus* | CM | CcCMc | 49.590912 | 22.639686 | SE | Primeval_protected |
| *Cucujus cinnaberinus* | CM | CcCMd | 49.433376 | 22.611067 | SE | Primeval_not_protected |
| *Cucujus cinnaberinus* | CM | CcCMe | 49.710492 | 22.731971 | SE | Primeval_not_protected |
| *Cucujus cinnaberinus* | CM | CcCMf | 49.683663 | 22.605898 | SE | Primeval_not_protected |
| *Cucujus cinnaberinus* | CM | CcCMg | 49.5765097 | 22.4696408 | SE | Primeval_protected |
| *Cucujus cinnaberinus* | DB | CcDBa | 51.53947 | 17.33627 | SW | Reserve |
| *Cucujus cinnaberinus* | DB | CcDBb | 51.53287 | 17.34944 | SW | Reserve |
| *Cucujus cinnaberinus* | OF | CcGOa | 49.93805 | 18.334733 | SW | Commercial |
| *Cucujus cinnaberinus* | OF | CcGOb | 50.010983 | 18.265917 | SW | Commercial |
| *Cucujus cinnaberinus* | OF | CcDOa | 52.392222 | 14.540556 | SW | Reserve |
| *Cucujus cinnaberinus* | OF | CcDOb | 52.367348 | 14.560756 | SW | Reserve |
| *Boros schneideri* | BF | BsBFa | 52.69084 | 23.727883 | NE | Primeval_not_protected |
| *Boros schneideri* | BF | BsBFb | 52.691053 | 23.728781 | NE | Primeval_not_protected |
| *Boros schneideri* | BF | BsBFc | 52.668435 | 23.768888 | NE | Primeval_not_protected |
| *Boros schneideri* | BF | BsBFd | 52.676158 | 23.846566 | NE | Primeval_not_protected |
| *Boros schneideri* | BF | BsBFe | 52.689022 | 23.794917 | NE | Primeval_not_protected |
| *Boros schneideri* | BF | BsBFf | 52.633086 | 23.717552 | NE | Primeval_protected |
| *Boros schneideri* | BF | BsBFg | 52.6802 | 23.75 | NE | Primeval_not_protected |
| *Boros schneideri* | BF | BsBFh | 52.6777 | 23.7513 | NE | Primeval_not_protected |
| *Boros schneideri* | KF | BsKFa | 53.2641 | 23.215724 | NE | Commercial |
| *Boros schneideri* | KF | BsKFb | 53.063729 | 23.727651 | NE | Commercial |
| *Boros schneideri* | KF | BsKFc | 53.148504 | 23.681801 | NE | Commercial |
| *Boros schneideri* | KF | BsKFd | 53.147472 | 23.679471 | NE | Commercial |
| *Boros schneideri* | KF | BsKFe | 53.146083 | 23.681364 | NE | Commercial |
| *Boros schneideri* | KF | BsKFf | 53.1544 | 23.8226 | NE | Commercial |
| *Boros schneideri* | AF | BsAFa | 53.86137 | 23.330817 | NE | Reserve |
| *Boros schneideri* | AF | BsAFb | 53.870625 | 23.299634 | NE | Reserve |
| *Boros schneideri* | AF | BsAFc | 53.872536 | 23.305556 | NE | Reserve |
| *Boros schneideri* | AF | BsAFd | 53.876493 | 23.296686 | NE | Reserve |
| *Boros schneideri* | AF | BsAFe | 53.91163 | 23.233723 | NE | Commercial |
| *Boros schneideri* | AF | BsAFf | 53.88132 | 23.379145 | NE | Commercial |
| *Boros schneideri* | AF | BsAFg | 53.850222 | 23.306545 | NE | Commercial |
| *Boros schneideri* | AF | BsAFh | 53.99144 | 23.477051 | NE | Commercial |
| *Boros schneideri* | AF | BsAFi | 53.900467 | 23.416372 | NE | Commercial |
| *Boros schneideri* | HM | BsHMa | 51.06814 | 20.721668 | SE | Primeval_protected |
| *Boros schneideri* | HM | BsHMb | 51.075652 | 20.765236 | SE | Primeval_protected |
| *Boros schneideri* | HM | BsHMc | 51.069231 | 20.733275 | SE | Primeval_protected |
| *Boros schneideri* | HM | BsHMd | 51.081609 | 20.763396 | SE | Primeval_protected |
| *Boros schneideri* | HM | BsHMe | 51.043391 | 20.683144 | SE | Primeval_protected |
| *Boros schneideri* | CM | BsCMa | 49.53679 | 22.630264 | SE | Primeval_protected |
| *Boros schneideri* | CM | BsCMb | 49.59744 | 22.621848 | SE | Primeval_protected |
| *Boros schneideri* | CM | BsCMc | 49.433069 | 22.611171 | SE | Primeval_protected |
| *Boros schneideri* | CM | BsCMd | 49.430055 | 22.620355 | SE | Primeval_protected |
| *Boros schneideri* | CM | BsCMe | 49.433672 | 22.63153 | SE | Primeval_protected |
| *Boros schneideri* | CM | BsCMf | 49.601293 | 22.673111 | SE | Primeval_protected |
| *Elater ferrugineus* | BF | EfBFa | 52.848657 | 23.877893 | NE | Primeval_not_protected |
| *Elater ferrugineus* | BF | EfBFb | 52.817592 | 23.759258 | NE | Primeval_protected |
| *Elater ferrugineus* | BF | EfBFc | 52.833065 | 23.683877 | NE | Primeval_not_protected |
| *Elater ferrugineus* | BF | EfBFd | 52.654272 | 23.65124 | NE | Primeval_protected |
| *Elater ferrugineus* | BF | EfBFe | 52.640601 | 23.697837 | NE | Primeval_protected |
| *Elater ferrugineus* | BF | EfBFf | 52.6522 | 23.7734 | NE | Primeval_protected |
| *Elater ferrugineus* | BF | EfBFg | 52.7419 | 23.7762 | NE | Primeval_not_protected |
| *Elater ferrugineus* | BF | EfBFh | 52.6932 | 23.8137 | NE | Primeval_protected |
| *Elater ferrugineus* | BF | EfBFi | 52.7698 | 23.7599 | NE | Primeval_protected |
| *Elater ferrugineus* | HM | EfHMa | 51.059055 | 20.701547 | SE | Primeval_protected |
| *Elater ferrugineus* | HM | EfHMb | 50.926786 | 20.919799 | SE | Primeval_protected |
| *Elater ferrugineus* | HM | EfHMc | 50.880654 | 21.103255 | SE | Primeval_protected |
| *Elater ferrugineus* | DB | EfDBa | 51.50814 | 17.02761 | SW | Reserve |
| *Elater ferrugineus* | DB | EfDBb | 51.5366 | 17.38831 | SW | Reserve |
| *Elater ferrugineus* | DB | EfDBc | 51.5366 | 17.38831 | SW | Reserve |
| *Elater ferrugineus* | DB | EfDBd | 51.51619 | 17.02687 | SW | Commercial |
| *Elater ferrugineus* | DB | EfDBe | 51.51619 | 17.02687 | SW | Commercial |
| *Elater ferrugineus* | DB | EfDBf | 51.47857 | 16.92432 | SW | Commercial |
| *Elater ferrugineus* | DB | EfDBg | 51.47857 | 16.92432 | SW | Commercial |
| *Elater ferrugineus* | DB | EfDBh | 51.47857 | 16.92432 | SW | Commercial |
| *Elater ferrugineus* | DB | EfDBi | 51.47857 | 16.92432 | SW | Commercial |
| *Elater ferrugineus* | DB | EfDBj | 51.59687 | 17.24105 | SW | Commercial |
| *Elater ferrugineus* | DB | EfDBk | 51.59687 | 17.24105 | SW | Commercial |
| *Elater ferrugineus* | DB | EfDBl | 51.47467 | 17.38537 | SW | Commercial |
| *Elater ferrugineus* | DB | EfDBm | 51.47467 | 17.38537 | SW | Commercial |
| *Elater ferrugineus* | DB | EfDBn | 51.47648126 | 17.01709148 | SW | Commercial |
| *Elater ferrugineus* | DB | EfDBo | 51.532206 | 17.335939 | SW | Commercial |
| *Elater ferrugineus* | OF | EfSFa | 50.946325 | 17.328137 | SW | Reserve |
| *Elater ferrugineus* | OF | EfSFb | 50.942147 | 17.390499 | SW | Reserve |
| *Elater ferrugineus* | OF | EfSFc | 50.942147 | 17.390499 | SW | Reserve |
| *Elater ferrugineus* | SF | EfBDa | 51.50753 | 15.70049 | SW | Reserve |
| *Elater ferrugineus* | SF | EfBDb | 51.50574 | 15.69694 | SW | Reserve |
| *Elater ferrugineus* | SF | EfBDc | 51.5019984 | 15.7038027 | SW | Commercial |
| *Elater ferrugineus* | SF | EfBDd | 51.5234 | 15.76449 | SW | Reserve |
| *Elater ferrugineus* | SF | EfBDe | 51.3431676 | 15.4322648 | SW | Commercial |
| *Elater ferrugineus* | SF | EfBDf | 51.34233 | 15.43239 | SW | Commercial |
| *Elater ferrugineus* | SF | EfBDg | 51.39802 | 15.46055 | SW | Commercial |
| *Elater ferrugineus* | SF | EfBDh | 51.33381 | 15.42478 | SW | Commercial |
| *Neomida haemorrhoidalis* | BF | NhBFa | 52.87256 | 23.627128 | NE | Primeval_not_protected |
| *Neomida haemorrhoidalis* | BF | NhBFb | 52.876691 | 23.61505 | NE | Primeval_not_protected |
| *Neomida haemorrhoidalis* | BF | NhBFc | 52.631994 | 23.692121 | NE | Primeval_protected |
| *Neomida haemorrhoidalis* | BF | NhBFd | 52.619985 | 23.721947 | NE | Primeval_protected |
| *Neomida haemorrhoidalis* | BF | NhBFe | 52.650353 | 23.76027 | NE | Primeval_not_protected |
| *Neomida haemorrhoidalis* | BF | NhBFf | 52.786346 | 23.712682 | NE | Primeval_not_protected |
| *Neomida haemorrhoidalis* | KF | NhKFa | 53.207705 | 23.581453 | NE | Reserve |
| *Neomida haemorrhoidalis* | KF | NhKFb | 53.338764 | 23.30416 | NE | Reserve |
| *Neomida haemorrhoidalis* | KF | NhKFc | 53.287037 | 23.110281 | NE | Reserve |
| *Neomida haemorrhoidalis* | KF | NhKFd | 53.285427 | 23.108097 | NE | Reserve |
| *Neomida haemorrhoidalis* | KF | NhKFe | 53.290852 | 23.29906 | NE | Commercial |
| *Neomida haemorrhoidalis* | AF | NhAFa | 54.026576 | 23.505948 | NE | Reserve |
| *Neomida haemorrhoidalis* | AF | NhAFb | 53.870315 | 23.305854 | NE | Reserve |
| *Neomida haemorrhoidalis* | AF | NhAFc | 53.838159 | 23.425035 | NE | Commercial |
| *Neomida haemorrhoidalis* | AF | NhAFd | 53.926458 | 23.424194 | NE | Commercial |
| *Neomida haemorrhoidalis* | AF | NhAFe | 53.901598 | 23.139493 | NE | Commercial |
| *Neomida haemorrhoidalis* | HM | NhHMa | 50.886943 | 21.105477 | SE | Primeval_protected |
| *Neomida haemorrhoidalis* | HM | NhHMb | 50.885282 | 21.105302 | SE | Primeval_protected |
| *Neomida haemorrhoidalis* | HM | NhHMc | 50.887996 | 21.097949 | SE | Primeval_protected |
| *Neomida haemorrhoidalis* | HM | NhHMd | 51.059055 | 20.701547 | SE | Primeval_protected |
| *Neomida haemorrhoidalis* | HM | NhHMe | 51.058089 | 20.706233 | SE | Primeval_protected |
| *Neomida haemorrhoidalis* | HM | NhHMf | 51.054532 | 20.705351 | SE | Primeval_protected |
| *Neomida haemorrhoidalis* | CM | NhCMa | 49.518117 | 22.419412 | SE | Primeval_protected |
| *Neomida haemorrhoidalis* | CM | NhCMb | 49.515292 | 22.425648 | SE | Primeval_protected |
| *Neomida haemorrhoidalis* | CM | NhCMc | 49.53254 | 22.568425 | SE | Primeval_protected |
| *Neomida haemorrhoidalis* | CM | NhCMd | 49.527903 | 22.573531 | SE | Primeval_protected |
| *Neomida haemorrhoidalis* | CM | NhCMe | 49.567374 | 22.587728 | SE | Primeval_protected |
| *Neomida haemorrhoidalis* | CM | NhCMf | 49.563968 | 22.590637 | SE | Primeval_protected |
| *Neomida haemorrhoidalis* | SF | NhBDa | 51.533611 | 17.513042 | SW | Commercial |
| *Neomida haemorrhoidalis* | SF | NhBDb | 51.5372 | 17.39519 | SW | Reserve |
| *Neomida haemorrhoidalis* | SF | NhBDc | 51.505917 | 16.93765 | SW | Commercial |
| *Neomida haemorrhoidalis* | SF | NhBDd | 51.467833 | 17.264483 | SW | Reserve |
| *Neomida haemorrhoidalis* | SF | NhBDe | 51.440533 | 17.249117 | SW | Commercial |
| *Neomida haemorrhoidalis* | SF | NhBDf | 51.440533 | 17.249117 | SW | Commercial |
| *Neomida haemorrhoidalis* | DB | NhDBa | 51.49745 | 15.694467 | SW | Reserve |
| *Neomida haemorrhoidalis* | DB | NhDBb | 51.501417 | 15.694617 | SW | Reserve |
| *Neomida haemorrhoidalis* | DB | NhDBc | 51.503217 | 15.691617 | SW | Reserve |
| *Neomida haemorrhoidalis* | DB | NhDBd | 51.505533 | 15.684717 | SW | Reserve |
| *Neomida haemorrhoidalis* | DB | NhDBe | 51.508567 | 15.67905 | SW | Reserve |
| *Neomida haemorrhoidalis* | DB | NhDBf | 51.50145 | 15.684567 | SW | Commercial |
| *Bolitophagus reticulatus* | BF | BrBFa | 52.704049 | 23.6943 | NE | Primeval_protected |
| *Bolitophagus reticulatus* | BF | BrBFb | 52.787066 | 23.796406 | NE | Primeval_not_protected |
| *Bolitophagus reticulatus* | BF | BrBFc | 52.87256 | 23.627128 | NE | Primeval_not_protected |
| *Bolitophagus reticulatus* | BF | BrBFd | 52.633414 | 23.728642 | NE | Primeval_not_protected |
| *Bolitophagus reticulatus* | BF | BrBFe | 52.7992 | 23.6435 | NE | Primeval_protected |
| *Bolitophagus reticulatus* | BF | BrBFf | 52.6143 | 23.5965 | NE | Primeval_not_protected |
| *Bolitophagus reticulatus* | KF | BrKFa | 53.276617 | 23.382295 | NE | Reserve |
| *Bolitophagus reticulatus* | KF | BrKFb | 53.207705 | 23.581453 | NE | Reserve |
| *Bolitophagus reticulatus* | KF | BrKFc | 53.338764 | 23.30416 | NE | Reserve |
| *Bolitophagus reticulatus* | KF | BrKFd | 53.287037 | 23.110281 | NE | Reserve |
| *Bolitophagus reticulatus* | KF | BrKFe | 53.33912 | 23.327452 | NE | Commercial |
| *Bolitophagus reticulatus* | KF | BrKFf | 53.401518 | 23.319261 | NE | Commercial |
| *Bolitophagus reticulatus* | KF | BrKFg | 53.345095 | 23.23897 | NE | Commercial |
| *Bolitophagus reticulatus* | AF | BrAFa | 54.043262 | 23.469799 | NE | Reserve |
| *Bolitophagus reticulatus* | AF | BrAFb | 54.026576 | 23.505948 | NE | Reserve |
| *Bolitophagus reticulatus* | AF | BrAFc | 53.880487 | 23.338363 | NE | Reserve |
| *Bolitophagus reticulatus* | AF | BrAFd | 54.06116 | 23.361667 | NE | Commercial |
| *Bolitophagus reticulatus* | AF | BrAFe | 53.926458 | 23.424194 | NE | Commercial |
| *Bolitophagus reticulatus* | AF | BrAFf | 53.901598 | 23.139493 | NE | Commercial |
| *Bolitophagus reticulatus* | HM | BrHMa | 50.966209 | 20.476825 | SE | Primeval_protected |
| *Bolitophagus reticulatus* | HM | BrHMb | 50.885282 | 21.105302 | SE | Primeval_not_protected |
| *Bolitophagus reticulatus* | HM | BrHMc | 50.886943 | 21.105477 | SE | Primeval_not_protected |
| *Bolitophagus reticulatus* | HM | BrHMd | 51.059055 | 20.701547 | SE | Primeval_protected |
| *Bolitophagus reticulatus* | HM | BrHMe | 51.058089 | 20.706233 | SE | Primeval_protected |
| *Bolitophagus reticulatus* | CM | BrCMa | 49.656216 | 22.49454 | SE | Primeval_protected |
| *Bolitophagus reticulatus* | CM | BrCMb | 49.572939 | 22.474125 | SE | Primeval_protected |
| *Bolitophagus reticulatus* | CM | BrCMc | 49.595515 | 22.633238 | SE | Primeval_protected |
| *Bolitophagus reticulatus* | CM | BrCMd | 49.532723 | 22.568631 | SE | Primeval_protected |
| *Bolitophagus reticulatus* | CM | BrCMe | 49.567045 | 22.584735 | SE | Primeval_protected |
| *Bolitophagus reticulatus* | CM | BrCMf | 49.612747 | 22.642828 | SE | Primeval_not_protected |
| *Bolitophagus reticulatus* | CM | BrCMg | 49.582607 | 22.471822 | SE | Primeval_not_protected |
| *Bolitophagus reticulatus* | CM | BrCMh | 49.557876 | 22.500507 | SE | Primeval_not_protected |
| *Bolitophagus reticulatus* | DB | BrDBa | 51.533352 | 17.512821 | SW | Commercial |
| *Bolitophagus reticulatus* | DB | BrDBb | 51.47372 | 17.41899 | SW | Commercial |
| *Bolitophagus reticulatus* | DB | BrDBc | 51.473581 | 17.423193 | SW | Commercial |
| *Bolitophagus reticulatus* | DB | BrDBd | 51.440533 | 17.249117 | SW | Commercial |
| *Bolitophagus reticulatus* | DB | BrDBe | 51.439278 | 17.255534 | SW | Commercial |
| *Bolitophagus reticulatus* | SF | BrSFa | 51.49745 | 15.694467 | SW | Reserve |
| *Bolitophagus reticulatus* | SF | BrSFb | 51.521121 | 15.772386 | SW | Reserve |
| *Schizotus pectinnicornis* | BF | SpBFa | 52.676158 | 23.846566 | NE | Primeval_not_protected |
| *Schizotus pectinnicornis* | BF | SpBFb | 52.6804 | 23.864762 | NE | Primeval_not_protected |
| *Schizotus pectinnicornis* | BF | SpBFc | 52.687247 | 23.8612 | NE | Primeval_not_protected |
| *Schizotus pectinnicornis* | BF | SpBFd | 52.702839 | 23.889652 | NE | Primeval_not_protected |
| *Schizotus pectinnicornis* | BF | SpBFe | 52.721709 | 23.916196 | NE | Primeval_not_protected |
| *Schizotus pectinnicornis* | KF | SpKFa | 53.410621 | 23.380008 | NE | Reserve |
| *Schizotus pectinnicornis* | KF | SpKFb | 53.415597 | 23.374172 | NE | Reserve |
| *Schizotus pectinnicornis* | KF | SpKFc | 53.342542 | 23.300179 | NE | Reserve |
| *Schizotus pectinnicornis* | KF | SpKFd | 53.276623 | 23.380327 | NE | Reserve |
| *Schizotus pectinnicornis* | KF | SpKFe | 53.2807 | 23.374351 | NE | Reserve |
| *Schizotus pectinnicornis* | KF | SpKFf | 53.114651 | 23.828271 | NE | Commercial |
| *Schizotus pectinnicornis* | KF | SpKFg | 53.111851 | 23.802333 | NE | Commercial |
| *Schizotus pectinnicornis* | AF | SpAFa | 54.043262 | 23.469799 | NE | Reserve |
| *Schizotus pectinnicornis* | AF | SpAFb | 53.853938 | 23.388729 | NE | Commercial |
| *Schizotus pectinnicornis* | AF | SpAFc | 53.838777 | 23.404268 | NE | Commercial |
| *Schizotus pectinnicornis* | AF | SpAFd | 54.06116 | 23.361667 | NE | Commercial |
| *Schizotus pectinnicornis* | AF | SpAFe | 54.060037 | 23.35966 | NE | Reserve |
| *Schizotus pectinnicornis* | HM | SpHMa | 50.884848 | 21.115949 | SE | Primeval_protected |
| *Schizotus pectinnicornis* | HM | SpHMb | 51.076384 | 20.717132 | SE | Primeval_not_protected |
| *Schizotus pectinnicornis* | HM | SpHMc | 51.077229 | 20.766369 | SE | Primeval_not_protected |
| *Schizotus pectinnicornis* | HM | SpHMd | 51.058996 | 20.70225 | SE | Primeval_protected |
| *Schizotus pectinnicornis* | HM | SpHMe | 51.0529 | 20.7027 | SE | Primeval_protected |
| *Schizotus pectinnicornis* | CM | SpCMa | 49.696139 | 22.538882 | SE | Primeval_protected |
| *Schizotus pectinnicornis* | CM | SpCMb | 49.538222 | 22.558827 | SE | Primeval_protected |
| *Schizotus pectinnicornis* | CM | SpCMc | 49.564666 | 22.594576 | SE | Primeval_protected |
| *Schizotus pectinnicornis* | CM | SpCMd | 49.591002 | 22.639542 | SE | Primeval_protected |
| *Schizotus pectinnicornis* | CM | SpCMe | 49.706319 | 22.693283 | SE | Primeval_not_protected |
| *Schizotus pectinnicornis* | SF | SpBDa | 51.503217 | 15.691617 | SW | Reserve |
| *Schizotus pectinnicornis* | SF | SpBDb | 51.505533 | 15.684717 | SW | Reserve |
| *Schizotus pectinnicornis* | DB | SpDBa | 51.53544 | 17.39263 | SW | Reserve |
| *Schizotus pectinnicornis* | DB | SpDBb | 51.50425 | 16.93835 | SW | Reserve |
| *Schizotus pectinnicornis* | DB | SpDBc | 51.52721 | 17.25745 | SW | Commercial |
| *Schizotus pectinnicornis* | DB | SpDBd | 51.472212 | 17.377078 | SW | Commercial |
| *Schizotus pectinnicornis* | DB | SpSFa | 51.533808 | 17.513715 | SW | Commercial |
| *Schizotus pectinnicornis* | OF | SpSFb | 49.93805 | 18.334733 | SW | Commercial |
| *Schizotus pectinnicornis* | OF | SpSFc | 50.013417 | 18.2726 | SW | Commercial |
| *Schizotus pectinnicornis* | OF | SpSFd | 51.04001 | 17.20826 | SW | Commercial |
| *Stenurella melanura* | BF | SmBFa | 52.7406 | 23.7679 | NE | Primeval_not_protected |
| *Stenurella melanura* | BF | SmBFb | 52.7457 | 23.772 | NE | Primeval_protected |
| *Stenurella melanura* | BF | SmBFc | 52.7153 | 23.6569 | NE | Primeval_protected |
| *Stenurella melanura* | BF | SmBFd | 52.6596 | 23.6211 | NE | Primeval_not_protected |
| *Stenurella melanura* | BF | SmBFe | 52.8455 | 23.8437 | NE | Primeval_not_protected |
| *Stenurella melanura* | KF | SmKFa | 53.3337 | 23.0622 | NE | Reserve |
| *Stenurella melanura* | KF | SmKFb | 53.3001 | 23.1263 | NE | Reserve |
| *Stenurella melanura* | KF | SmKFc | 53.291 | 23.1089 | NE | Reserve |
| *Stenurella melanura* | KF | SmKFd | 53.168198 | 23.839165 | NE | Commercial |
| *Stenurella melanura* | KF | SmKFe | 53.240492 | 23.745095 | NE | Commercial |
| *Stenurella melanura* | KF | SmKFf | 53.173406 | 23.789747 | NE | Commercial |
| *Stenurella melanura* | KF | SmKFg | 53.497999 | 23.319043 | NE | Commercial |
| *Stenurella melanura* | AF | SmAFa | 53.8964 | 23.3154 | NE | Reserve |
| *Stenurella melanura* | AF | SmAFb | 53.8727 | 23.3052 | NE | Reserve |
| *Stenurella melanura* | AF | SmAFc | 53.8741 | 23.3486 | NE | Reserve |
| *Stenurella melanura* | AF | SmAFd | 53.925743 | 23.231447 | NE | Commercial |
| *Stenurella melanura* | AF | SmAFe | 53.901091 | 23.218148 | NE | Commercial |
| *Stenurella melanura* | AF | SmAFf | 53.907562 | 23.3282 | NE | Commercial |
| *Stenurella melanura* | AF | SmAFg | 54.014246 | 23.349533 | NE | Commercial |
| *Stenurella melanura* | HM | SmHMa | 51.059055 | 20.701547 | SE | Primeval_protected |
| *Stenurella melanura* | HM | SmHMb | 50.962024 | 20.476074 | SE | Primeval_protected |
| *Stenurella melanura* | HM | SmHMc | 50.9197 | 20.9003 | SE | Primeval_protected |
| *Stenurella melanura* | HM | SmHMd | 50.927187 | 20.919478 | SE | Primeval_protected |
| *Stenurella melanura* | HM | SmHMe | 51.060741 | 20.696823 | SE | Primeval_not_protected |
| *Stenurella melanura* | CM | SmCMa | 49.799266 | 22.662891 | SE | Primeval_protected |
| *Stenurella melanura* | CM | SmCMb | 49.660891 | 22.509354 | SE | Primeval_protected |
| *Stenurella melanura* | CM | SmCMc | 49.694744 | 22.538115 | SE | Primeval_protected |
| *Stenurella melanura* | CM | SmCMd | 49.518067 | 22.418805 | SE | Primeval_protected |
| *Stenurella melanura* | CM | SmCMe | 49.456178 | 22.50602 | SE | Primeval_not_protected |
| *Stenurella melanura* | DB | SmDBa | 51.51957 | 15.76484 | SW | Reserve |
| *Stenurella melanura* | DB | SmDBb | 51.502898 | 16.997124 | SW | Reserve |
| *Stenurella melanura* | DB | SmDBc | 51.50796642 | 17.02785745 | SW | Reserve |
| *Stenurella melanura* | DB | SmDBd | 51.50117 | 17.34964 | SW | Commercial |
| *Stenurella melanura* | DB | SmDBe | 51.56313 | 16.84588 | SW | Commercial |
| *Stenurella melanura* | OF | SmSFa | 50.9327 | 17.409883 | SW | Commercial |
| *Stenurella melanura* | OF | SmSFb | 50.94335 | 17.392983 | SW | Reserve |
| *Stenurella melanura* | OF | SmSFc | 50.9275 | 17.40745 | SW | Commercial |
| *Stenurella melanura* | OF | SmSFd | 50.936917 | 17.394717 | SW | Commercial |
| *Stenurella melanura* | OF | SmSFe | 51.88055 | 15.754 | SW | Commercial |
| *Stenurella melanura* | SF | SmBDa | 51.50753 | 15.70049 | SW | Reserve |
| *Stenurella melanura* | SF | SmBDb | 51.34055 | 15.43079 | SW | Commercial |
| *Stenurella melanura* | SF | SmBDc | 51.38291 | 15.41588 | SW | Commercial |
| *Stenurella melanura* | SF | SmBDd | 51.502462 | 15.733386 | SW | Commercial |
| *Rhagium inquisitor* | BF | RiBFa | 52.6646 | 23.7307 | NE | Primeval_not_protected |
| *Rhagium inquisitor* | BF | RiBFb | 52.711239 | 23.923645 | NE | Primeval_not_protected |
| *Rhagium inquisitor* | BF | RiBFc | 52.6292 | 23.7131 | NE | Primeval_protected |
| *Rhagium inquisitor* | BF | RiBFd | 52.689022 | 23.794917 | NE | Primeval_not_protected |
| *Rhagium inquisitor* | BF | RiBFe | 52.854557 | 23.660515 | NE | Primeval_not_protected |
| *Rhagium inquisitor* | KF | RiKFa | 53.207705 | 23.581453 | NE | Reserve |
| *Rhagium inquisitor* | KF | RiKFb | 53.410621 | 23.380008 | NE | Reserve |
| *Rhagium inquisitor* | KF | RiKFc | 53.342542 | 23.300179 | NE | Reserve |
| *Rhagium inquisitor* | KF | RiKFd | 53.300037 | 23.126508 | NE | Reserve |
| *Rhagium inquisitor* | KF | RiKFe | 53.142032 | 23.676242 | NE | Commercial |
| *Rhagium inquisitor* | KF | RiKFf | 53.192781 | 23.668165 | NE | Commercial |
| *Rhagium inquisitor* | AF | RiAFa | 53.872536 | 23.305556 | NE | Reserve |
| *Rhagium inquisitor* | AF | RiAFb | 53.871591 | 23.348753 | NE | Reserve |
| *Rhagium inquisitor* | AF | RiAFc | 54.02699 | 23.480565 | NE | Reserve |
| *Rhagium inquisitor* | AF | RiAFd | 54.040475 | 23.470555 | NE | Reserve |
| *Rhagium inquisitor* | AF | RiAFe | 53.91163 | 23.233723 | NE | Commercial |
| *Rhagium inquisitor* | AF | RiAFf | 53.88132 | 23.379145 | NE | Commercial |
| *Rhagium inquisitor* | AF | RiAFg | 53.821271 | 23.339823 | NE | Commercial |
| *Rhagium inquisitor* | AF | RiAFh | 53.99144 | 23.477051 | NE | Commercial |
| *Rhagium inquisitor* | HM | RiHMa | 51.04342 | 20.755186 | SE | Primeval_not_protected |
| *Rhagium inquisitor* | HM | RiHMb | 50.886739 | 21.076081 | SE | Primeval_protected |
| *Rhagium inquisitor* | HM | RiHMc | 51.081756 | 20.772031 | SE | Primeval_not_protected |
| *Rhagium inquisitor* | HM | RiHMd | 50.966557 | 20.479139 | SE | Primeval_protected |
| *Rhagium inquisitor* | HM | RiHMe | 50.82551 | 21.113214 | SE | Primeval_not_protected |
| *Rhagium inquisitor* | CM | RiCMa | 49.528312 | 22.57142 | SE | Primeval_protected |
| *Rhagium inquisitor* | CM | RiCMb | 49.683928 | 22.60343 | SE | Primeval_not_protected |
| *Rhagium inquisitor* | CM | RiCMc | 49.433345 | 22.611151 | SE | Primeval_not_protected |
| *Rhagium inquisitor* | CM | RiCMd | 49.671528 | 22.479754 | SE | Primeval_not_protected |
| *Rhagium inquisitor* | CM | RiCMe | 49.706991 | 22.693458 | SE | Primeval_not_protected |
| *Rhagium inquisitor* | DB | RiDBa | 51.533202 | 17.561304 | SW | Commercial |
| *Rhagium inquisitor* | DB | RiDBb | 51.438467 | 17.22055 | SW | Commercial |
| *Rhagium inquisitor* | SF | RiSFa | 51.502533 | 15.731967 | SW | Commercial |
| *Rhagium inquisitor* | SF | RiSFb | 51.498483 | 15.713983 | SW | Commercial |
| *Rhagium inquisitor* | DB | RiSFc | 51.532433 | 17.093767 | SW | Reserve |
| *Thanasimus formicarius* | BF | TfBFa | 52.682857 | 23.729106 | NE | Primeval_not_protected |
| *Thanasimus formicarius* | BF | TfBFf | 52.682291 | 23.727025 | NE | Primeval_not_protected |
| *Thanasimus formicarius* | BF | TfBFb | 52.637575 | 23.549957 | NE | Primeval_not_protected |
| *Thanasimus formicarius* | BF | TfBFc | 52.7239 | 23.6729 | NE | Primeval_protected |
| *Thanasimus formicarius* | BF | TfBFd | 52.6663 | 23.7131 | NE | Primeval_not_protected |
| *Thanasimus formicarius* | BF | TfBFe | 52.696839 | 23.886439 | NE | Primeval_not_protected |
| *Thanasimus formicarius* | KF | TfKFa | 53.1481 | 23.6824 | NE | Commercial |
| *Thanasimus formicarius* | KF | TfKFb | 53.153973 | 23.888846 | NE | Commercial |
| *Thanasimus formicarius* | KF | TfKFc | 53.151427 | 23.821783 | NE | Commercial |
| *Thanasimus formicarius* | KF | TfKFd | 53.127971 | 23.822929 | NE | Commercial |
| *Thanasimus formicarius* | KF | TfKFe | 53.11717 | 23.613402 | NE | Commercial |
| *Thanasimus formicarius* | KF | TfKFf | 53.28903 | 23.134141 | NE | Commercial |
| *Thanasimus formicarius* | AF | TfAFa | 53.870625 | 23.299634 | NE | Reserve |
| *Thanasimus formicarius* | AF | TfAFb | 53.871853 | 23.298037 | NE | Reserve |
| *Thanasimus formicarius* | AF | TfAFc | 53.870625 | 23.299634 | NE | Reserve |
| *Thanasimus formicarius* | AF | TfAFd | 54.0403 | 23.4705 | NE | Reserve |
| *Thanasimus formicarius* | AF | TfAFe | 54.0403 | 23.4705 | NE | Reserve |
| *Thanasimus formicarius* | AF | TfAFf | 53.849094 | 23.364357 | NE | Commercial |
| *Thanasimus formicarius* | AF | TfAFg | 53.905572 | 23.327901 | NE | Commercial |
| *Thanasimus formicarius* | AF | TfAFh | 53.960924 | 23.480475 | NE | Commercial |
| *Thanasimus formicarius* | HM | TfHMa | 51.077229 | 20.766369 | SE | Primeval_not_protected |
| *Thanasimus formicarius* | HM | TfHMb | 51.076269 | 20.765329 | SE | Primeval_not_protected |
| *Thanasimus formicarius* | HM | TfHMc | 51.049725 | 20.680722 | SE | Primeval_not_protected |
| *Thanasimus formicarius* | HM | TfHMd | 51.053367 | 20.703742 | SE | Primeval_protected |
| *Thanasimus formicarius* | HM | TfHMe | 51.052741 | 20.707919 | SE | Primeval_not_protected |
| *Thanasimus formicarius* | CM | TfCMa | 49.422822 | 22.652495 | SE | Primeval_not_protected |
| *Thanasimus formicarius* | CM | TfCMb | 49.430367 | 22.648826 | SE | Primeval_not_protected |
| *Thanasimus formicarius* | CM | TfCMc | 49.661685 | 22.499918 | SE | Primeval_protected |
| *Thanasimus formicarius* | CM | TfCMd | 49.673638 | 22.455682 | SE | Primeval_not_protected |
| *Thanasimus formicarius* | CM | TfCMe | 49.641556 | 22.483103 | SE | Primeval_not_protected |
| *Thanasimus formicarius* | DB | TfDBa | 51.525311 | 17.338445 | SW | Commercial |
| *Thanasimus formicarius* | DB | TfDBb | 51.44725 | 17.261117 | SW | Commercial |
| *Thanasimus formicarius* | OF | TfLZa | 51.3129 | 16.422867 | SW | Commercial |
| *Thanasimus formicarius* | SF | TfSFa | 51.50107 | 15.69765 | SW | Reserve |
| *Thanasimus formicarius* | SF | TfSFb | 51.52051 | 15.76959 | SW | Commercial |
| *Thanasimus formicarius* | SF | TfSFc | 51.3274425 | 15.4429949 | SW | Commercial |
| *Thanasimus formicarius* | SF | TfSFd | 51.498856 | 15.908514 | SW | Commercial |
| *Thanasimus formicarius* | SF | TfGOa | 51.510917 | 15.757583 | SW | Commercial |
| *Phaenops cyanea* | BF | PcBFa | 52.60347 | 23.586405 | NE | Primeval_not_protected |
| *Phaenops cyanea* | BF | PcBFb | 52.615406 | 23.638547 | NE | Primeval_protected |
| *Phaenops cyanea* | BF | PcBFc | 52.645491 | 23.700817 | NE | Primeval_protected |
| *Phaenops cyanea* | BF | PcBFd | 52.64553 | 23.695867 | NE | Primeval_not_protected |
| *Phaenops cyanea* | KF | PcKFa | 53.1481 | 23.6824 | NE | Commercial |
| *Phaenops cyanea* | KF | PcKFb | 53.139647 | 23.740088 | NE | Commercial |
| *Phaenops cyanea* | KF | PcKFc | 53.138064 | 23.729059 | NE | Commercial |
| *Phaenops cyanea* | KF | PcKFd | 53.1382 | 23.4458 | NE | Commercial |
| *Phaenops cyanea* | KF | PcKFe | 53.139614 | 23.667175 | NE | Commercial |
| *Phaenops cyanea* | AF | PcAFa | 53.905572 | 23.327901 | NE | Commercial |
| *Phaenops cyanea* | AF | PcAFb | 53.827152 | 23.48394 | NE | Commercial |
| *Phaenops cyanea* | AF | PcAFc | 53.828678 | 23.485109 | NE | Commercial |
| *Phaenops cyanea* | AF | PcAFd | 53.858552 | 23.329734 | NE | Commercial |
| *Phaenops cyanea* | AF | PcAFe | 54.0117 | 23.4922 | NE | Commercial |
| *Phaenops cyanea* | HM | PcHMa | 50.8553 | 20.5904 | SE | Primeval_not_protected |
| *Phaenops cyanea* | HM | PcHMb | 51.072171 | 20.759632 | SE | Primeval_not_protected |
| *Phaenops cyanea* | HM | PcHMc | 51.077229 | 20.766369 | SE | Primeval_not_protected |
| *Phaenops cyanea* | CM | PcCMa | 49.645911 | 22.505886 | SE | Primeval_not_protected |
| *Phaenops cyanea* | CM | PcCMb | 49.674069 | 22.450204 | SE | Primeval_not_protected |
| *Phaenops cyanea* | CM | PcCMc | 49.641401 | 22.483742 | SE | Primeval_not_protected |
| *Phaenops cyanea* | CM | PcCMd | 49.604684 | 22.586708 | SE | Primeval_not_protected |
| *Phaenops cyanea* | CM | PcCMe | 49.689106 | 22.409715 | SE | Primeval_not_protected |
| *Phaenops cyanea* | CM | PcCMf | 49.723196 | 22.605758 | SE | Primeval_not_protected |
| *Phaenops cyanea* | DB | PcDBa | 51.54507 | 17.02616 | SW | Commercial |
| *Phaenops cyanea* | SF | PcSFa | 51.391244 | 15.5394047 | SW | Commercial |
| *Phaenops cyanea* | SF | PcSFb | 51.51323 | 15.75438 | SW | Commercial |
| *Phaenops cyanea* | SF | PcSFc | 51.50063 | 15.7214 | SW | Commercial |
| *Phaenops cyanea* | SF | PcSFd | 51.4178 | 15.85139 | SW | Commercial |
| *Ips typographus* | BF | ItBFa | 52.711239 | 23.923645 | NE | Primeval_not_protected |
| *Ips typographus* | BF | ItBFb | 52.615198 | 23.638882 | NE | Primeval_protected |
| *Ips typographus* | BF | ItBFc | 52.792468 | 23.79715 | NE | Primeval_not_protected |
| *Ips typographus* | BF | ItBFd | 52.678664 | 23.683029 | NE | Primeval_protected |
| *Ips typographus* | BF | ItBFe | 52.7239 | 23.6729 | NE | Primeval_protected |
| *Ips typographus* | BF | ItBFf | 52.7216 | 23.7469 | NE | Primeval_not_protected |
| *Ips typographus* | BF | ItBFg | 52.719677 | 23.747305 | NE | Primeval_protected |
| *Ips typographus* | BF | ItBFh | 52.637575 | 23.549957 | NE | Primeval_not_protected |
| *Ips typographus* | KF | ItKFa | 53.207705 | 23.581453 | NE | Reserve |
| *Ips typographus* | KF | ItKFb | 53.205956 | 23.580292 | NE | Reserve |
| *Ips typographus* | KF | ItKFc | 53.345631 | 23.313411 | NE | Reserve |
| *Ips typographus* | KF | ItKFd | 53.2807 | 23.374351 | NE | Reserve |
| *Ips typographus* | KF | ItKFe | 53.27746 | 23.115688 | NE | Reserve |
| *Ips typographus* | KF | ItKFf | 53.300037 | 23.126508 | NE | Reserve |
| *Ips typographus* | KF | ItKFg | 53.137386 | 23.717442 | NE | Commercial |
| *Ips typographus* | KF | ItKFh | 53.1481 | 23.6824 | NE | Commercial |
| *Ips typographus* | KF | ItKFi | 53.147295 | 23.679492 | NE | Commercial |
| *Ips typographus* | KF | ItKFj | 53.121627 | 23.860565 | NE | Commercial |
| *Ips typographus* | KF | ItKFk | 53.28903 | 23.134141 | NE | Commercial |
| *Ips typographus* | AF | ItAFa | 53.870625 | 23.299634 | NE | Reserve |
| *Ips typographus* | AF | ItAFb | 53.870038 | 23.303833 | NE | Reserve |
| *Ips typographus* | AF | ItAFc | 53.871853 | 23.298037 | NE | Reserve |
| *Ips typographus* | AF | ItAFd | 53.849094 | 23.364357 | NE | Commercial |
| *Ips typographus* | AF | ItAFe | 53.99144 | 23.477051 | NE | Commercial |
| *Ips typographus* | AF | ItAFf | 53.905572 | 23.327901 | NE | Commercial |
| *Ips typographus* | AF | ItAFg | 53.932207 | 23.439236 | NE | Commercial |
| *Ips typographus* | AF | ItAFh | 53.959978 | 23.510733 | NE | Commercial |
| *Ips typographus* | AF | ItAFi | 53.827152 | 23.48394 | NE | Commercial |
| *Ips typographus* | AF | ItAFj | 53.901091 | 23.218148 | NE | Commercial |
| *Ips typographus* | HM | ItHMa | 51.068395 | 20.738438 | SE | Primeval_protected |
| *Ips typographus* | HM | ItHMb | 51.046542 | 20.669862 | SE | Primeval_not_protected |
| *Ips typographus* | HM | ItHMc | 51.086704 | 20.72645 | SE | Primeval_not_protected |
| *Ips typographus* | CM | ItCMa | 49.284948 | 22.683412 | SE | Primeval_not_protected |
| *Ips typographus* | CM | ItCMb | 49.423119 | 22.652127 | SE | Primeval_not_protected |
| *Ips typographus* | CM | ItCMc | 49.430454 | 22.648068 | SE | Primeval_not_protected |
| *Ips typographus* | CM | ItCMd | 49.430454 | 22.648068 | SE | Primeval_not_protected |
| *Ips typographus* | CM | ItCMe | 49.430454 | 22.648068 | SE | Primeval_not_protected |
| *Ips typographus* | DB | DB-It | 51.489542 | 17.118366 | SW | Commercial |
| *Ips typographus* | DB | DB-It | 51.487905 | 17.118251 | SW | Commercial |
| *Ips typographus* | OF | GO-It | 50.97671 | 17.382455 | SW | Commercial |
| *Ips typographus* | OF | GO-It | 51.395771 | 16.463509 | SW | Commercial |
| *Ips typographus* | OF | GO-It | 51.285543 | 16.876407 | SW | Commercial |
| *Ips typographus* | OF | GO-It | 51.009955 | 17.398202 | SW | Commercial |
| *Ips typographus* | SF | It-SF | 51.339137 | 15.261584 | SW | Commercial |
| *Ips acuminatus* | BF | IaBFa | 52.6383 | 23.5499 | NE | Primeval_not_protected |
| *Ips acuminatus* | BF | IaBFb | 52.723 | 23.6073 | NE | Primeval_protected |
| *Ips acuminatus* | BF | IaBFc | 52.6865 | 23.7162 | NE | Primeval_not_protected |
| *Ips acuminatus* | BF | IaBFd | 52.6775 | 23.8097 | NE | Primeval_not_protected |
| *Ips acuminatus* | KF | IaKFa | 53.1451 | 23.797 | NE | Commercial |
| *Ips acuminatus* | KF | IaKFb | 53.1489 | 23.7933 | NE | Commercial |
| *Ips acuminatus* | KF | IaKFc | 53.1274 | 23.3386 | NE | Commercial |
| *Ips acuminatus* | KF | IaKFd | 53.2802 | 23.3564 | NE | Commercial |
| *Ips acuminatus* | KF | IaKFe | 53.1481 | 23.6824 | NE | Commercial |
| *Ips acuminatus* | KF | IaKFf | 53.151427 | 23.821783 | NE | Commercial |
| *Ips acuminatus* | AF | IaAFa | 53.9007 | 23.2199 | NE | Commercial |
| *Ips acuminatus* | AF | IaAFb | 53.9255 | 23.2311 | NE | Commercial |
| *Ips acuminatus* | AF | IaAFc | 53.9195 | 23.235 | NE | Commercial |
| *Ips acuminatus* | AF | IaAFd | 53.8966 | 23.3318 | NE | Commercial |
| *Ips acuminatus* | AF | IaAFe | 53.9333 | 23.3149 | NE | Commercial |
| *Ips acuminatus* | AF | IaAFf | 53.9598 | 23.5108 | NE | Commercial |
| *Ips acuminatus* | AF | IaAFg | 53.8866 | 23.2276 | NE | Commercial |
| *Ips acuminatus* | HM | IaHMa | 51.075652 | 20.765236 | SE | Primeval_not_protected |
| *Ips acuminatus* | HM | IaHMb | 50.885605 | 21.078882 | SE | Primeval_protected |
| *Ips acuminatus* | HM | IaHMc | 50.885807 | 21.071652 | SE | Primeval_protected |
| *Ips acuminatus* | HM | IaHMd | 50.867651 | 21.063475 | SE | Primeval_protected |
| *Ips acuminatus* | OF | IaSFa | 51.000324 | 17.447994 | SW | Commercial |
| *Ips acuminatus* | OF | IaSFb | 50.983289 | 17.377978 | SW | Commercial |
| *Ips acuminatus* | OF | IaSFc | 51.010703 | 17.399337 | SW | Commercial |
| *Ips acuminatus* | OF | IaSFd | 50.789988 | 18.441084 | SW | Commercial |
| *Ips acuminatus* | OF | IaSFe | 51.011946 | 17.39893 | SW | Commercial |
| *Ips acuminatus* | SF | IaBDa | 51.339137 | 15.261584 | SW | Commercial |
| *Ips acuminatus* | SF | IaBDb | 51.326283 | 15.496797 | SW | Commercial |

Table S2: Sample sizes (number of collected beetles) across the experimental groups, based on geography regions (NE: north east, SE: south east, SW: south west) and based on management level (C: commercial, R: reserve, Pn: primeval not protected, Pp: primeval protected).

| Species | No. of Ind. | No. of SNPs | No. of loci | No. of SNPs (LD-pruned) | Geography | | | Management | | | | |
| --- | --- | --- | --- | --- | --- | --- | --- | --- | --- | --- | --- | --- |
|  |  |  |  |  | NE | SE | SW | | C | R | Pn | Pp |
| *Neomida haemorrhoidalis* | 199 | 10501 | 4553 | 7378 | 79 | 60 | 60 | | 35 | 74 | 20 | 69 |
| *Boros schneideri* | 154 | 6574 | 2649 | 4639 | 97 | 57 | 0 | | 49 | 24 | 45 | 40 |
| *Cucujus cinnaberinus* | 190 | 2480 | 1336 | 1943 | 103 | 59 | 28 | | 60 | 50 | 41 | 39 |
| *Elater ferrugineus* | 84 | 10210 | 4184 | 4788 | 40 | 10 | 34 | | 20 | 14 | 15 | 35 |
| *Bolitophagus reticulatus* | 185 | 2841 | 1460 | 1973 | 90 | 61 | 34 | | 54 | 45 | 39 | 47 |
| *Stenurella melanura* | 199 | 12063 | 2882 | 8069 | 93 | 47 | 57 | | 73 | 53 | 23 | 48 |
| *Schizotus pectinicornis* | 167 | 2512 | 1107 | 1835 | 80 | 47 | 40 | | 45 | 51 | 38 | 33 |
| *Thanasimus formicarius* | 178 | 2426 | 434 | 951 | 99 | 41 | 38 | | 78 | 30 | 60 | 10 |
| *Ips acuminatus* | 125 | 3957 | 1517 | 2507 | 84 | 20 | 21 | | 85 | 0 | 20 | 20 |
| *Ips typographus* | 201 | 1484 | 568 | 1049 | 132 | 34 | 35 | | 87 | 40 | 54 | 20 |
| *Rhagium inquisitor* | 164 | 4083 | 1144 | 2668 | 95 | 46 | 23 | | 48 | 40 | 57 | 19 |
| *Phaenops cyanea* | 140 | 27583 | 9321 | 16590 | 70 | 45 | 25 | | 75 | 0 | 55 | 10 |
